# How useful is genomic data for predicting maladaptation to future climate?

**DOI:** 10.1101/2023.02.10.528022

**Authors:** Brandon M. Lind, Rafael Candido-Ribeiro, Pooja Singh, Mengmeng Lu, Dragana Obreht Vidakovic, Tom R. Booker, Michael C. Whitlock, Sam Yeaman, Nathalie Isabel, Sally N. Aitken

## Abstract

Methods using genomic information to forecast potential population maladaptation to climate change are becoming increasingly common, yet the lack of model validation poses serious hurdles toward their incorporation into management and policy. Here, we compare the validation of maladaptation estimates derived from two methods – Gradient Forests (GF_offset_) and the Risk Of Non-Adaptedness (RONA) – using exome capture pool-seq data from 35 to 39 populations across three conifer taxa: two Douglas-fir varieties and jack pine. We evaluate sensitivity of these algorithms to the source of input loci (markers selected from genotype-environment associations [GEA] or those selected at random). We validate these methods against two-year and 52-year growth and mortality measured in independent transplant experiments. Overall, we find that both methods often better predict transplant performance than climatic or geographic distances. We also find that GF_offset_ and RONA models are surprisingly not improved using GEA candidates. Even with promising validation results, variation in model projections to future climates makes it difficult to identify the most maladapted populations using either method. Our work advances understanding of the sensitivity and applicability of these approaches, and we discuss recommendations for their future use.

## 1 Introduction

Environmental and land use change pose unprecedented risk to global biodiversity loss (Exposito-Alonso et al., 2022; Nadeau et al., 2017; Urban, 2015). Historically, the impacts of these changes on species’ distributions have been projected through species distribution modelling (e.g., Thuiller et al., 2008). However, these methods often fail to account for the environmental drivers of local adaptation or the various evolutionary mechanisms (e.g., geno flow, phenotypic plasticity) by which populations could respond to environmental change (ONeill et al., 2008; Waldvogel et al., 2020). Recently, methods incorporating genomic information to forecast climate maladaptation have gained traction. Predominant among these, Gradient Forests (GF_offset_, *sensu* Fitzpatrick & Keller, 2015) and the Risk Of Non-Adaptedness (RONA; Rellstab et al., 2016) use current relationships between genotype and climate to estimate genomic offset (i.e., a measure of maladaptation between a population’s current or future environment and its environmental optimum; see Methods). Often used to estimate maladaptation to future climate change, these offset methods can also incorporate non-climatic environmental variables. If such offset methods were shown to be robust when estimating maladaptation to any future environmental factor, they could circumvent the need for long-term field experiments and could rapidly inform management priorities, or provide an option for species where experimentation is not feasible.

Despite their current popularity, genomic offset methods remain largely unvalidated with a few exceptions. For instance, (Láruson et al., 2022) used simulated data to evaluate GF_offset_. They found that when 1) climate and genotypes are known without error, 2) all populations across the simulated landscape are locally adapted, and 3) validation is carried out within the climate space used in training, the predicted offset had a strong negative rank correlation with simulated fitness, and GF_offset_ models trained using all markers performed no better than GF_offset_ models trained using causal markers. Further, they found that environmental distances calculated using environmental variables driving local adaptation also had a strong negative relationship with simulated fitness, though this was not the case when non-causal environments were included in distance calculations.

Compared to simulated data, attempts to validate GF_offset_ using empirical data where error is inherent have found relatively weaker relationships between GF_offset_ and juvenile phenotypes measured in a common garden (Fitzpatrick et al., 2021). This suggests that offset models may not perform as well in practice as they do under ideal circumstances. Even so, and similar to findings of Láruson et al. (2022), Fitzpatrick et al. (2021) also found GF_offset_ to be more accurate than naïve climate distances, further suggesting genomic offset methods of this and other types (e.g., Capblancq & Forester, 2021) provide advantages over climate data alone. However, the relatively weaker empirical performance than that found from simulation data may be improved with more direct measures of survival and reproduction.

In empirical settings, GF_offset_ models are often used to project offset to areas of the species’ range where no populations have been sampled and to climates many decades into the future (e.g., Bay et al., 2018; Fitzpatrick & Keller, 2015; Gougherty et al., 2021; Lu et al., 2019; Vanhove et al., 2021) . However, projection to unsampled environments can lead to inaccuracies when models cannot generalize well. While generalizability poses one hurdle, it is unclear whether more accessible forms of data (e.g., climate or geographic distance) perform as well as these genetically based methods in all systems. Further, offset implementations have used disparate sets and sample sizes of both populations and loci to project future offset to changing climates, without exploring the impact of these sources on model predictions (but see e.g., Fitzpatrick et al., 2021; Láruson et al., 2022).

Validating these methods’ predictions of maladaptation to future climate is challenging due to the temporal nature of such projections. However, transplant experiments (i.e., common gardens or provenance trials) can be used to quantify performance by correlating measurements of fitness-related phenotypes with the offset projected to the contemporary climate of the growing site (Blois et al., 2013; Fitzpatrick et al., 2021).

Tree species are ideally suited to empirically validate predictions from offset methods because there is abundant evidence from transplant experiments to suggest that many tree species are locally adapted to climate (Boshier et al., 2015; Lind et al., 2018; Savolainen et al., 2007; Sork et al., 2013), a key underlying assumption of offset models (Capblancq et al., 2020; Láruson et al., 2022; Rellstab et al., 2021). Trees are also relevant systems for understanding maladaptation to future climate because of their ecological role in terrestrial systems, as well as their capacity to sequester carbon. Many forest tree species have experienced large geographic range shifts in the past in response to changes in climate (Davis & Shaw, 2001; Hamrick, 2004). Yet, rates of projected climate change are likely to outpace maximum rates of historical migration for many of these species (e.g., Dauphin et al., 2021; Davis & Shaw, 2001; McLachlan et al., 2005) and therefore leave future outcomes largely unknown (Alberto et al., 2013; Allen et al., 2010; Mckenney et al., 2007; Millar et al., 2007).

Here, we train offset models using exome capture pool-seq data from three conifer taxa (Fig. 1) and validate results with phenotypes from independent transplant experiments at seedling (two-year Douglas-fir, *Pseudotsuga menziesii*) and adult (52-year jack pine, *Pinus banksiana*) life stages. Using fitness-related phenotypes from juvenile life stages enables validation of projections for species where no longer-term phenotypic data exist (the situation for most species), while validation using phenotypes from adult life stages enables comparison of offset to more direct measures of total lifetime fitness. The main goal of this study is to use empirical datasets for widespread species known to be locally adapted to climate to evaluate potential consequences of decisions made during training and validation. Specifically, we use these datasets to address four main questions: Q1) How is performance of the offset method affected by the source of training loci? Q2) How do genomic offsets compare with non-genomic offset measures of climate and geographic distance? Q3) How are inferences from models affected by the populations used for validation?

**Fig. 1.**
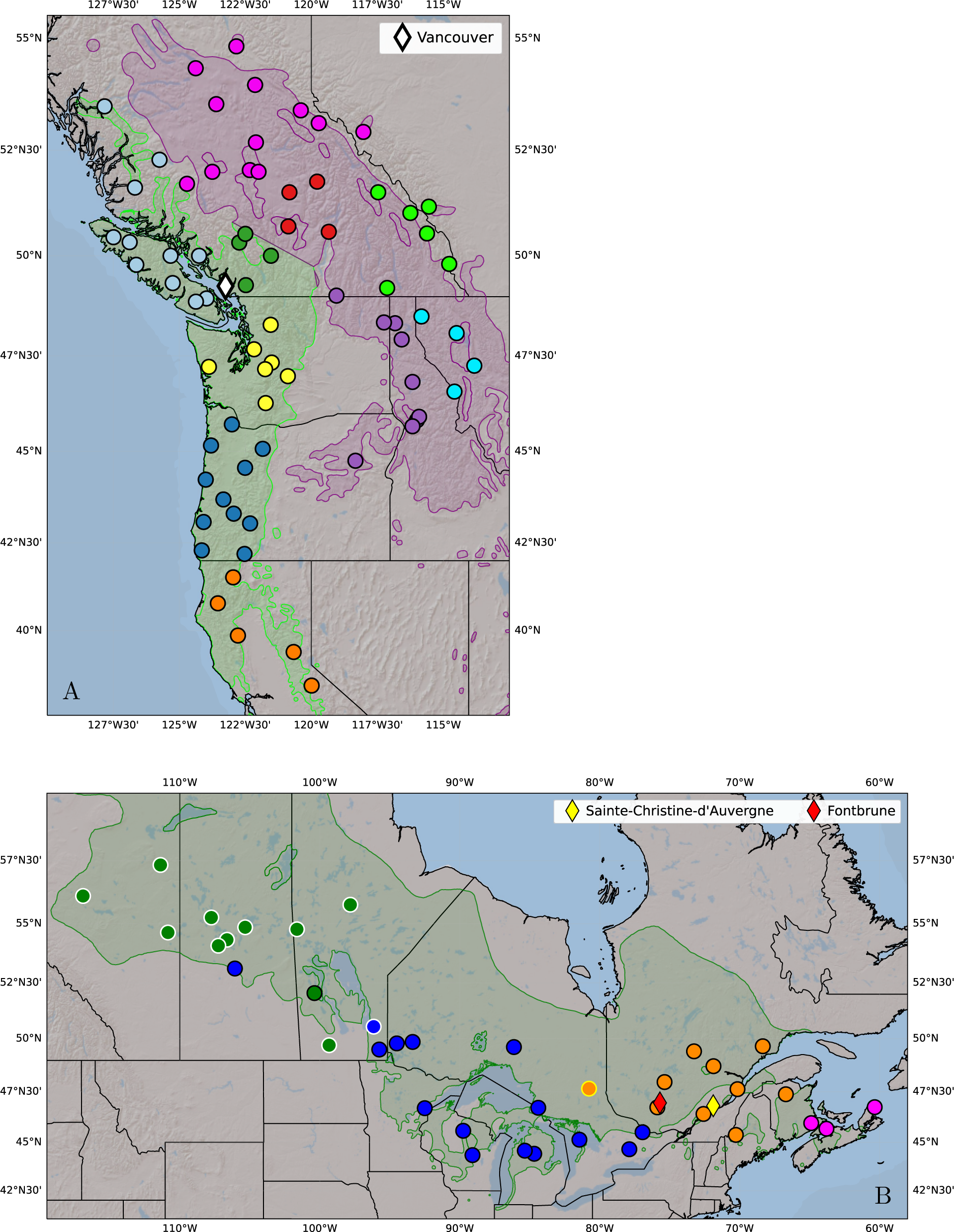
North American source populations (circles) used for genomic and phenotypic data to train and validate both Gradient Forests (GF) and the Risk of Non-Adaptedness (RONA), and the common gardens used for offset validation: (A) Douglas-fir, and (B) jack pine. Shaded polygons are range maps for the full range of coastal Douglas-fir (lime, A), the northern and central range of interior Douglas-fir (purple, A), and the southern range of jack pine (green, B). All Douglas-fir populations were grown in the Vancouver common garden (white diamond, A) and used for validation. Jack pine populations outlined in black were used for validation in both the Sainte-Christine (yellow diamond, B) and Fontbrune (red diamond, B) common gardens, while those outlined in yellow were only used for validation in Sainte-Christine, and those outlined in white were only used in model training but not validation. Color of population indicates the genetic groups used for visualization. Code to generate these figures can be found in SN 15.99.

## 2 Methods

Throughout this manuscript we will be referencing our code used to carry out specific analyses in-line with the text, most often in unstripped jupyter notebooks (Kluyver et al., 2016). We refer to these notebooks as Supplemental Notebooks (SN) using a directory numbering system (e.g., SN 15.01). More information about the numbering system and archiving can be found in the Data Availability section.

### 2.1 Focal Species, Population Sampling, and Genetic Data

Three taxa of conifers across two species (Fig. 1) were used to assess the accuracy of genomic offset methods: 1) 38 range-wide populations of coastal Douglas-fir (*Pseudotsuga menziesii* var. *menziesii* [Mirb.] Franco, *Pinaceae*), 2) 35 populations of interior Douglas-fir (*P*. *menziesii* var. *glauca*) from the variety’s northern and central range, and 3) 39 jack pine (*Pinus banksiana* Lamb., *Pinaceae*) populations from across the species’ southern range. We chose these species for their large and environmentally heterogeneous distributions, economic importance, and ecological relevance. Further, there is extensive evidence for local adaptation to climate in both varieties of Douglas-fir (Bansal, Clair, et al., 2015; Bansal, Harrington, et al., 2015; Krueger & Ferrell, 1965; Rehfeldt et al., 2014) as well as jack pine (Eckert et al., 2012; Rehfeldt et al., 1999, 2001; Wang, Hamann, et al., 2006; Wu & Ying, 2004).

We used exome capture pool-seq data from these sampled populations. Briefly, we targeted exomic regions in DNA extracted from 33 to 40 individuals per population for Douglas-fir (39 Mb capture probe size), and 17 to 20 individuals per population for jack pine (41 Mb capture probe size) using exome capture probes described in (Lind et al., 2022), where individuals within populations were pooled in equimolar quantities before sequencing. The sequencing depths used here exceed those used in (Lind et al., 2022), as it was found that pool-seq depth was one of the best predictors of the agreement of allele frequencies between sequence data for individuals and those from pool-seq data, despite generally strong agreement overall (Pearson’s r > 0.948; Lind et al., 2022).

Pool-seq libraries were sequenced in a 150bp paired-end format on an Illumina HiSeq4000 instrument at the Centre d’expertise et de Services Génome Québec, Montréal, Canada. We mapped reads from both varieties of Douglas-fir to the current reference genome of coastal Douglas-fir v1.01 (Neale et al., 2017). We mapped reads from jack pine to an amended version of its congener, loblolly pine (Neale et al., 2014; Wegrzyn et al., 2014; Zimin et al., 2014) (*P. taeda* L., Pinaceae v2.01). In short, we used transcriptomic data from jack pine and amended non-mapping transcripts to the loblolly reference before mapping pool-seq data.

Single nucleotide polymorphisms (SNPs) were called independently for Douglas-fir and jack pine according to bioinformatic best practices as detailed in (Lind et al., 2022) using a VarScan pipeline (Lind, 2021) and filtered for missing data across populations ≤ 25%, minimum read depth per population per locus ≥ 8, and global minor allele frequency ≥ 0.05. Paralogs can lead to erroneous SNP calls due to misalignment to a reference genome (McKinney et al., 2017; Rellstab et al., 2019). As in Lind et al. (2022), these loci were also filtered with the VarScan pipeline using SNPs called from haploid megagametophyte tissue (see Lind et al. 2022 for more details). We also created a fourth ‘cross-variety’ SNP set by combining the unfiltered data from both varieties and applying the same filtering process as for the single variety datasets such as read depth, missing data, and MAF (SN 02.01.01).

To address Q1, we use two methods for identifying genotype-environment association (GEA) candidates to ensure that genomic offset performance was not solely the outcome of the chosen method, as well as random sets of loci with numbers matching those of candidate sets to ensure that the source of loci was also not affecting the outcome. BayPass (Gautier, 2015) is a single-locus GEA that evaluates support for each SNP independently for each environmental variable. We also use GEA results from the Weighted Z Analysis (WZA, Booker et al., 2023). The WZA uses information across closely linked loci within genomic windows (here genic regions) to assess GEA support at the window level for a given environmental variable. We performed GEA analyses using BayPass and WZA at the variety level for Douglas-fir (see SN subfolder 02.02) and the species level for jack pine (see SN subfolder 07.02). For BayPass, we identified all SNPs across all 19 climatic variables (Table S2) with Bayes Factor (BF) in deciban units (dB) ≥ 15 following Jeffrey’s rule indicating, at minimum, very strong support (Extended Data Table 1; see Supplemental Note 1.9 for more details about BayPass implementation). From the WZA output, we identified the top 500 genes associated with each of the 19 climatic variables using *p*-values from the WZA. From within these gene windows, we kept only those SNPs that had a Kendall’s τ ≥ 0.5, which was calculated by correlating the population-level allele frequencies with environmental values for each locus (Extended Data Table 1). When using the ‘cross-variety’ SNPs filtered jointly across both Douglas-fir varieties, we used the intersection of loci between 1) those that passed cross-variety filtering and 2) those that were also GEA hits within varieties (i.e., we did not perform GEA across both varieties together). Hereafter, the BayPass and WZA marker sets are also referred to more generally as ‘candidate’ marker sets.

Because we are interested in knowing how the input loci would affect genomic offset methods (Q1), we also created a ‘random’ set of loci of equal sample size as each of the two candidate sets by randomly choosing loci across our full datasets (SN 15.04). In total, we generated four sets of SNPs for each of the three conifer taxa to use in training (Extended Data Table 1). The comparison of models using random and candidate sets addresses questions related to criteria of input loci and its impact on model performance, and comparison among models using the random sets of loci address questions related to the impact of the number of input loci to model performance. For the main text we present results using marker sets from BayPass, WZA, and the set of random loci with same sample size as WZA, and present all sets together within Supplemental Information. We used sets of random markers in this way because of computational constraints, as opposed to creating many sets of random markers sampled with replacement (see e.g., Fitzpatrick et al. 2021). For estimating RONA (Rellstab et al., 2016), we used a subset of each of these marker sets so that only loci with significant linear models were included in RONA calculations (see Section 2.3, Extended Data Table 1).

### 2.2 Training and Predicting Offset with Gradient Forests

Gradient Forests is a machine learning algorithm that incorporates Random Forest ensemble learning by using climate to split nodes of allele frequencies for a given locus in a forest of decision trees, and uses this splitting information to construct monotonic turnover functions which are in turn aggregated and used to predict offset to future climate (Fitzpatrick & Keller, 2015). Random Forest ensembles are known to handle correlated features (e.g., environmental variables) without causing overfitting (Géron, 2022; Raschka & Mirjalili, 2019), and (Láruson et al., 2022) found that correlated features did not reduce performance of GF_offset_ models or misidentify causal environments.

We used candidate and random marker sets (Section 2.1) to train GF_offset_ (SN 15.04; Ellis et al., 2012; Smith et al., 2012) in R (v3.5.1; R Core Team 2018). For each marker set, we created training sets that included all available populations (Extended Data Figure 3A-B).(e.g., Borrell et al., 2019; Gougherty et al., 2021; Gugger et al., 2021; Vanhove et al., 2021)

The climate data used in training included climate normals from 19 climatic environmental variables between the years 1961-1990 predating much of the recent anthropogenic warming, downloaded from AdaptWest.com on Februrary 5, 2021 (AdaptWest-Project, 2021); AdaptWest data is generated using ClimateNA (Wang et al., 2016). These climate variables include those related to annual temperature (MAT, MWMT, MCMT, TD), 30-year minimum (EMT) and maximum (EXT) temperature extremes, annual precipitation (MAP, AHM, Eref, CMD), and the seasonality of both temperature (DD0, DD5, NFFD, FFP, bFFP, eFFP) and precipitation (MSP, SHM, PAS; see Table S2 for climatic abbreviations and units). These variables were selected *a priori* based upon relevance to the species’ biology and environmental variation across the species’ ranges. After clipping AdaptWest climate data (SN 15.03) to our species ranges (SN 15.02) using range maps from the United States Geological Survey (Little, 1971), training sets and training scripts (SN 15.05) were used to train models of GF_offset_ (SN 15.04 section 5). Each trained GF_offset_ model was used to predict offset to the climate of one (Douglas-fir) or two (jack pine) common gardens (SN 15.07) using the script created in SN 15.05 and using the default linear extrapolation. The climate data used for offset prediction (SN 15.06) was the average climate (obtained from ClimateNA GUI between July 2-9 2021, Wang et al., 2016; Table S2), of the common garden over the years in which the individuals were grown (see Section 2.5), and was treated as the novel (i.e., ‘future’) climate of each population in offset projections. Thirty-year extreme variables, such as EMT and EXT, were also averaged across the values given for the years grown in the common garden.

An added utility of GF is that it can identify climatic variables driving variation in genetic data, without the need to project offset to future climates, particularly if offset is not the primary goal. Gradient Forests outputs ranked environmental importance after being trained. This has shown promise in identifying environmental drivers underlying selection when using simulated data (Láruson et al., 2022), even when there are multiple correlated environmental variables. Using the candidate and random marker sets, we explore the consistency of environmental importance ranks. We also explore the consistency of environmental importance ranks between candidate and random marker sets that used all populations in training (SN 15.13). We found that GF was relatively insensitive to marker and population input with regard to environmental importance (Supplemental Text S1.3; Figs. S3-S7).

### 2.3 Estimating the Risk of Non-Adaptedness (RONA)

In addition to GF_offset_, we also used RONA (Rellstab et al., 2016) to estimate genomic offset. The offset estimated by RONA relies on linear relationships between allele frequencies for candidate loci and climatic variables. This estimation is carried out in four steps: 1) identifying candidate loci putatively underlying adaptation to the environment (e.g., from GEA), 2) subsetting this list for loci that also have significant linear models relating allele frequency with environmental variables, 3) using the current model of the linear relationship between population allele frequencies and environment to estimate the allele frequency for a single population in a new environment (e.g., a value from projected climate change or a common garden), and 4) averaging the absolute difference between current and estimated future allele frequencies across loci for a given population for a given climatic variable (see equation and Fig. 2 on p. 5913 of Rellstab et al., 2016).

**Fig. 2.**
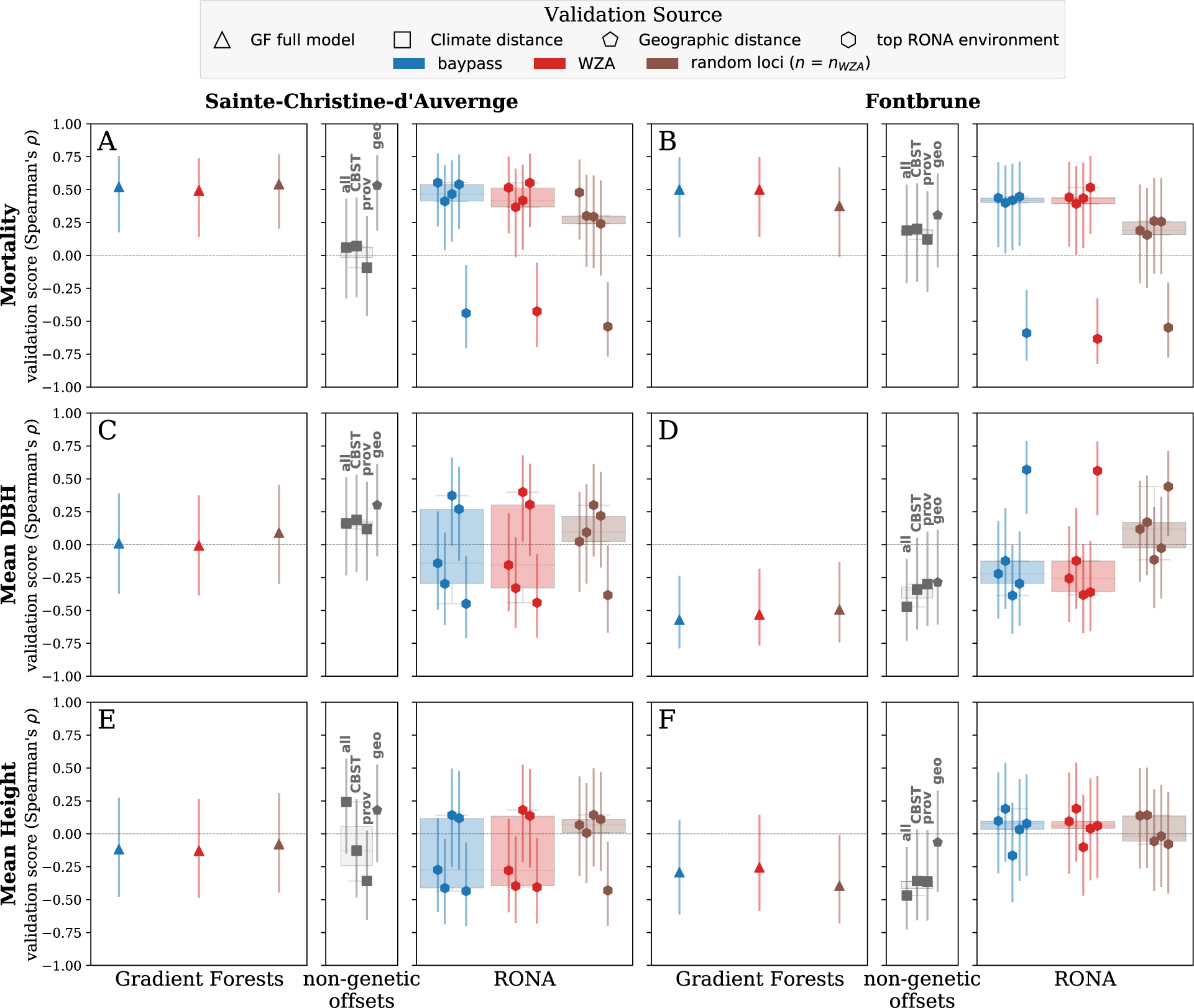
Offset validation from 52-year jack pine phenotypes at Sainte-Christine-d’Auvergne (A, C, E) and Fontbrune (B, D, F) provenance trials using Gradient Forests (GF_offset_), the Risk Of Non-Adaptedness (RONA), and climate and geographic distances. Triangles indicate performance of GF_offset_ models trained and validated using all available populations. RONA background boxplots illustrate the range of RONA validation scores given for the top five environmental variables (hexagons) that differed significantly between source and common garden variables (see Table S1). Climate distances (squares) were calculated using 1) all climate variables, or 2) those variables used for climate-based seed transfer (CBST) in British Columbia, or 3) those explaining significant variation in provenance trials. Vertical bars indicate standard error estimated using a Fisher transformation (see Supplemental Text S1.3). Loci used in RONA calculations are a subset of those used in GF_offset_ that had significant linear models with the environment, see Extended Data Table 1 for loci counts. See Fig. S8 for all locus groups. Boxplot whiskers extend up to 1.5x the interquartile range. See Extended Data Fig. 3 for a conceptual representation of training and validation sources. Code to create these figures can be found in SN 15.14.

Using the four marker sets described in Section 2.1 (two candidate sets and two random sets; Extended Data Figure 3A), we isolated loci with significant linear models relating current allele frequencies to climate variables (the same climate data used in GF_offset_ training in Section 2.2; Table S2), then calculated RONA for each population and environmental variable (SN 15.09) using average environmental values for the years individuals were grown in the gardens (Section 2.5). We grouped population-level predictions from the same population training sets used for GF_offset_ (Extended Data Figure 3B). Because RONA is calculated for a specific population and environmental variable, there is a range of RONA estimates for any given population, and thus the choice of environmental variables to consider for offset estimation could impact inferences regarding population performance in novel environments. To address this, Rellstab et al. (2016) used paired *t*-tests to determine which future environments were most different from their current state (*n* = 5), taking the top three variables after ranking *p*-values to use in estimating the range of RONA. For our validation, the vector containing future environments (i.e., common gardens) would be constant for a given variable across populations. In the context of a paired *t*-test, this is somewhat intractable with the test’s null hypotheses that each vector in the pair is sampled from the same distribution, which could lead to biologically meaningless (yet statistically significant) inference. We explored groups of environmental variables (see next section) related to ‘expert choice’ or those used in guiding seed sourcing in British Columbia. However, the top five environments from the original paired *t*-test as described above produced more accurate results (i.e., correct sign and higher magnitudes of Spearman’s *ρ*) than any of the other groups of environments (not shown, except in SN 15.09 section 8), and so we present the range of RONA using these top five environments from ranked *t*-test *p*-values. Because the top environments isolated in this way are often highly correlated with each other, these top environments will also likely give correlated offset estimates and thus allow for the effective estimation of one RONA offset value.

### 2.4 Non-genomic Offset Measures

Could environmental data alone be used instead of genetic data for management decisions (Mahony et al., 2020)? To compare genomic offsets with non-genomic offset measures of climate and geographic distance (Q2), we also estimated population offset by calculating geographic and climatic distances from the source populations to the common garden (SN 15.08). To calculate geographic distance, we use the latitude and longitude of each population and garden to calculate distance via Vincenty’s geodesic. To calculate climatic distance, we use the Mahalanobis distance for each population centered on the common garden using the same climate data in training and prediction with GF_offset_ and RONA (Sections 2.2 and 2.3; Table S2). We explored three sets of environmental variables to estimate climate distance: 1) all geoclimatic variables (Table S2), 2) those climate variables used in climate-based seed transfer (CBST) guidelines for British Columbia (O’Neill et al., 2009) – mean annual temperature (MAT), mean coldest month temperature (MCMT), continentality (TD), mean annual precipitation (MAP), degree-days above 5°C (DD5), extreme minimum temperature (EMT), and 3) climate variables identified in previous and independent reciprocal transplants not used here. For jack pine, we used the two climate variables from the transfer function used to best predict height of a sister species with which it readily hybridizes, lodgepole pine (*Pinus contorta* subsp. *latifolia* Douglas, *Pinaceae*)(Wang, Hanann, et al., 2006): MAT (>64% variance explained) and annual heat-moisture index (AHM; where *ln*(AHM) explained >6% variance). For Douglas-fir, we used three variables found to be significant predictors in universal response functions of height and basal diameter for a large multiple common garden trial of North American populations from both varieties planted in Central Europe (Chakraborty et al., 2015): MAT, summer heat-moisture index (SHM), and TD.

### 2.5 Common Garden Data

Measurements of fitness-related phenotypes from common gardens (diamonds, Fig. 1) used to validate genomic offset predictions were obtained by phenotyping individuals from the same populations that were genotyped. For jack pine, we measured 52-year adult phenotypes for height, diameter at breast height (DBH), and mortality in a field provenance trial at two sites, Fontbrune (LAT 46.959, LONG -75.698) and Sainte-Christine-d’Auvergne (LAT 46.819, LONG -71.888), between 1966 and 2018. For Douglas-fir, we measured two-year seedling phenotypes – shoot biomass and height increment – grown in a Vancouver common garden (LAT 49.257, LONG -123.250) between 2018-2019 (Candido-Ribeiro et al., 2022). For each common garden, we used the population mean phenotype to validate genomic offset (Section 2.6). For more information about phenotypic measurements, see Supplemental Text S1.4.

### 2.6 Validating Offset Measures

Population mean phenotypes (Section 2.5) were used as a proxy for fitness by which to validate the genomic offsets predicted from GF_offset_ (SN 15.11) and RONA (SN 15.09), by correlating population mean phenotype with population offset, using Spearman’s *ρ* as a validation score (Supplemental Text S1.5). Spearman’s *ρ* was used because we do not necessarily expect linear relationships between offset and phenotypes and wanted to explicitly test offsets in their ability to rank climate maladaptation, particularly given that offset and phenotypes are not measured in the same units (Lotterhos et al., 2022). If genomic offset is a good proxy for potential maladaptation, we expect a negative relationship between offset and growth, and a positive relationship between offset and mortality. For GF_offset_ models, we used all available offset estimates and phenotypes to calculate the validation score. We validated RONA for each environmental variable that ranked within the top five environments that differed significantly (via *t*-test *p*-values) between the common garden and climates used in training, calculating a validation score for each climate variable.

To determine if inference related to model performance was affected by the populations used in validation (Q3), we leveraged genetic structure within and across the two varieties of Douglas-fir (Extended Data Figure 3C). These two varieties (coastal and interior; green and purple ranges, respectively, in Fig. 1A) diverged ∼2.11 Ma (Gugger et al., 2010) and differ substantially both morphologically and ecologically. While the coastal variety shows little genetic grouping in PCA and instead differentiates along a latitudinal cline, populations in the northern range of interior Douglas-fir populations form two distinct genetic groups (Fig. S1). This allowed us to address Q3 by calculating our validation score using various levels of genetic hierarchy (Extended Data Fig. 3) – we used the offset predicted for either all or a subset of training populations to calculate validation scores. Specifically, we calculated validation scores across 1) populations from both varieties, 2) all interior variety populations, and 3) across populations from each of the northwestern and southeastern interior Douglas-fir genetic subgroups (see Fig. S1). We evaluate these hierarchical scenarios using the GF_offset_ models trained across both varieties as well as those trained using solely the interior variety (SN 15.11).

### 2.7 Projecting genomic offset to future climates

As in Section 2.2, we downloaded future climate scenarios from AdaptWest.com (AdaptWest-Project, 2021; Wang et al., 2016). We used GF_offset_ and RONA models trained with all WZA loci to project future genomic offsets to future climate scenarios (SN 15.07 and SN 15.16, respectively). For future climate scenarios we used Representative Concentration Pathway (RCP) greenhouse concentration trajectories projected to the 2050s and 2080s: RCP4.5 2050s, RCP4.5 2080s, RCP8.5 2050s, and RCP8.5 2080s. RCP4.5 and RCP8.5 each represent radiative forcing units (*W*/*m*^2^and are, respectively, an intermediate scenario where emissions peak in the 2040s and then decline, or continue to rise throughout the 21^st^ Century. As with estimating RONA using common gardens (Section 2.3), we identified the five environmental variables for which our sample populations differed the most between present and future climate scenarios. We report results from RCP8.5 2050s in the main text, including Spearman’s *ρ* between RONA estimates (SN 15.16), GF_offset_ (SN 15.18), and between estimates from both RONA and GF_offset_ (SN 15.16).

## 3 Results

### 3.1 Validation of offset with fitness-related phenotypes

#### 3.1.1 Jack pine

The performance of both GF_offset_ and RONA differed between the two jack pine provenance trials validated using 52-year phenotypes of mean DBH, mean height, and mortality (Fig. 2). Mortality often was better predicted than DBH and height. Genomic offset predictions of 52-year mortality were not demonstrably better than those based on the best non-genomic offset measure at either location. (Q2). Importantly, using candidate loci from GEA analyses did not improve predictive ability over randomly chosen loci for GF_offset_ (Q1; Fig. 2). For RONA the validation scores from GEA sets tended to have similar scores as random loci when estimating DBH and height, but scores from the two sets became more differentiated when estimating mortality (Q3; Fig. 2, Fig. S8).

The best non-genomic offset measure varied by phenotype and site, with low variation among validation scores for these metrics (Fig. 2). While geographic distance performed better than climate distances for mortality (Fig. 2A-B), climate distance tended to perform better for DBH and height (Fig. 2C-F), but the set of climate variables used to calculate the best distance varied, and only once exceeded the scores from the full GF_offset_ models (Fig. 2E).

#### 3.1.2 Douglas-fir

As with jack pine, genomic offsets estimated using random loci performed equally well as GEA sets (Q1). Both the cross-variety and coastal variety models from GF_offset_ and RONA substantially outperformed climate and geographic distance metrics for Douglas-fir, though this was not the case for the interior variety (Q2, Fig. 3, Extended Data Fig. 1). The GF_offset_ and RONA models that were trained and validated across both varieties of Douglas-fir had the greatest validation scores across all comparisons (Fig. 3A, Extended Data Fig. 1A), achieving much higher performance than in jack pine (Fig. 2). However, when models were trained and validated for each variety separately the relative performance decreased (Fig. 3B-C, Extended Data Fig. 1B-C). The stronger validation score from the cross-variety model validated using both varieties (e.g., Fig. 3A) compared to the scores validated within varieties is likely driven by the substantial genetic structure of the two varieties, as varieties are distinct when plotting cross-variety offset vs. phenotype (Extended Data Fig. 2).

**Fig. 3.**
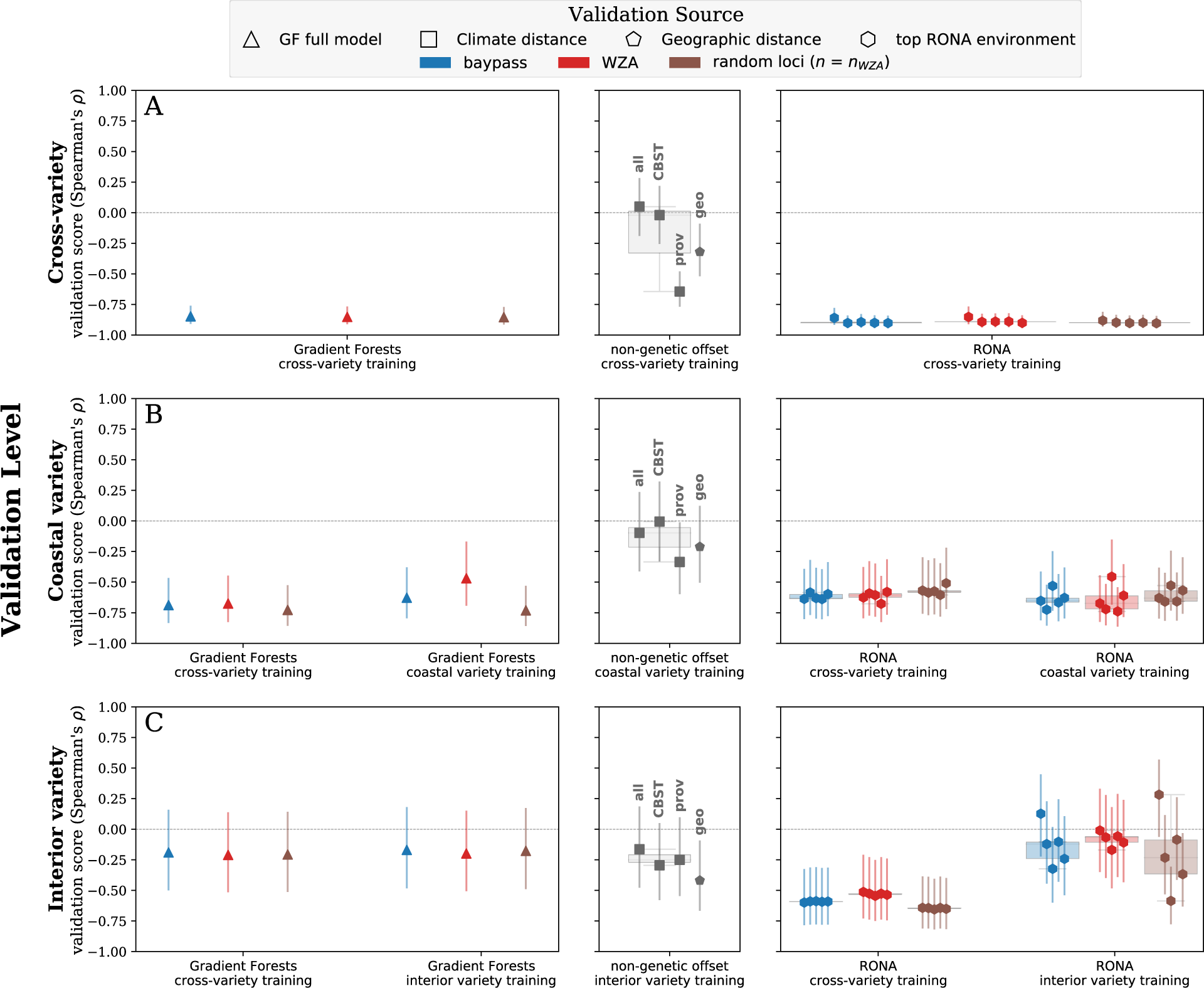
Offset validation from two-year Douglas-fir height increment phenotypes at the Vancouver common garden (see Fig. 1A) using Gradient Forests (GF_offset_), the Risk of Non-Adaptedness (RONA), and climate and geographic distances. We assessed accuracy inference from trained models (x-axis groups) using populations (rows) across both varieties of Douglas-fir (A), at the variety level for the coastal (B) and interior varieties of Douglas-fir (C) to determine if greater numbers of training populations improve finer-scale predictions of offset. Genetic offset boxplots and shapes are shaded with respect to marker set source. Triangles indicate performance of GF_offset_ models trained and validated using all available populations. RONA background boxplots illustrate the range of RONA validation scores given for the top five climatic variables (hexagons) that differed significantly between source population and the common garden (see Table S1). Climate distances (squares) were calculated using 1) all climate variables, or 2) those variables used for climate-based seed transfer (CBST) in British Columbia, or 3) those explaining significant variation in provenance trials. Vertical bars indicate standard error estimated using a Fisher transformation (see Supplemental Text S1.3). Loci used in RONA calculations are a subset of those used in Gradient Forests that had significant linear models with the environment, see Extended Data Table 1 for locus counts. See Extended Data Fig. 1 for similar validation using shoot biomass. See Fig. S9 for all locus groups. Boxplot whiskers extend up to 1.5x the interquartile range. See Extended Data Fig. 3 for a conceptual representation of training and validation sources. Code to create these figures can be found in SN 15.14.

Because management decisions are usually made at finer spatial scales than a species’ range, we were interested in how well groups of Douglas-fir populations (i.e., varieties or genetic groups) would validate, and if performance across all populations was indicative of performance at these finer spatial scales (Q3). Assessing performance at finer scales and with fewer populations than used in model training is particularly relevant. For instance, genetic structure in the data could lead to magnitudes of Spearman’s rho estimates that could be misinterpreted as a well-performing model, when in fact the model is a poor predictor at scales of management relevance (see Supplemental Text S1.10 for a toy example). Comparing models, the cross-variety model validated using only variety-specific populations was not substantially different from models that were both trained and validated at the variety level (Fig. 3B-C, Extended Data Fig. 1B-C). Comparing the two varieties, the coastal variety models had greater validation scores than models for the interior variety (Fig. 3B-C, Extended Data Fig. 1B-C). Coastal variety genomic offsets often performed better than non-genomic offset measures, but genomic and non-genomic offsets performed similarly for the interior variety (Q2, center panels Fig. 3, Extended Data Fig. 1). To further explore impacts on the accuracy of fine-scale offset, we subset populations from the interior variety into two distinct genetic groups to validate predictions from the GF_offset_ cross-variety and interior-only models. We found similar patterns of accuracy between fine-scale validation of the cross-variety and interior-only genomic offset models, though fine-scale validation indicated stronger relationships between offset and performance within these genetic groups than at the variety level (Supplemental Text S1.6).

Validation scores from climate distance using variables inferred as important from independent provenance trials were often stronger than the other climate distance measures (Fig. 3, Extended Data Fig. 1), while validation scores from geographic distance were stronger than climate distance only in interior Douglas-fir populations (Figs. S9-S10).

### 3.2 Predicted genomic offset to future climates

#### 3.2.1 Jack pine

Maladaptation of jack pine populations to future climate (RCP8.5 2050s) inferred from GF_offset_ and RONA models trained using WZA loci and all populations indicate that the western-most group (green populations, Fig. 1B) relative to all other populations are likely to experience the greatest maladaptive effects from changing climates (Fig. S11B-C).

These populations have consistently high maladaptive rank across both GF_offset_ and RONA (Fig. 4). From the projection of GF_offset_ to areas of the jack pine range with no training data, it would seem that the central portion of the range will be similarly maladapted to future climate (red contours, Fig. S12D). Across the five environmental variables used to estimate RONA for this climate scenario (which were highly correlated, Fig. S11E-F), the predicted maladaptive rank from RONA was positively correlated with GF_offset_ (Fig. S11C-D).

**Fig. 4.**
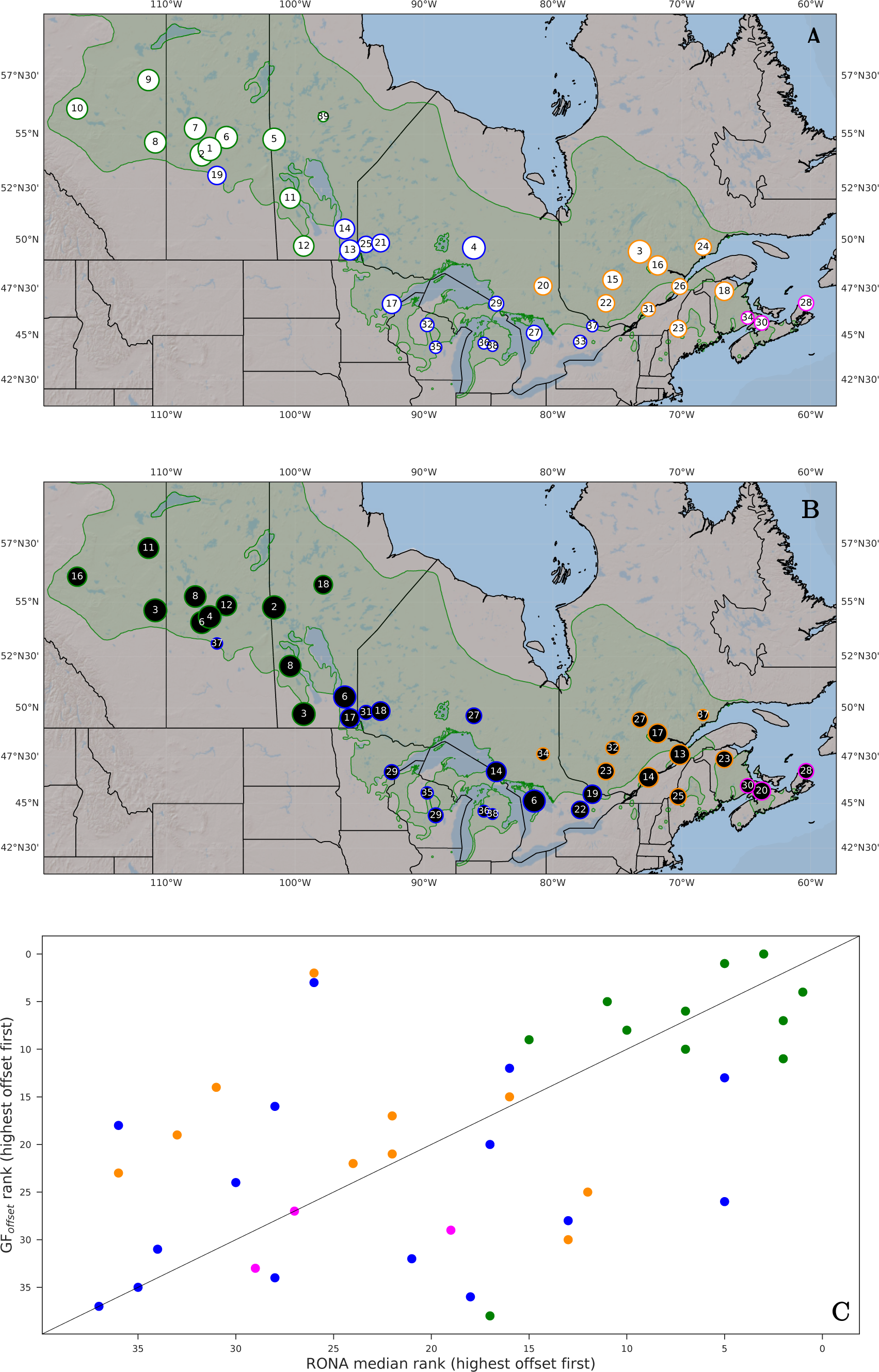
Maladaptation of jack pine populations to future climate (RCP8.5 2050s) inferred from Gradient Forests (GF_offset_, A, C) and RONA (B, C). Population point sizes in A and B are scaled to offset rank (lowest offset have smallest sizes). Population point sizes in B are from the median ranks across environments used to estimate RONA, which were chosen based on ranking p-values from paired *t-*tests between current and future climate. In C, a 1:1 line is given to infer relative changes in rank between methods. Rank numbers are given within circles of A and B. Colors correspond to groups of Fig. 1. Code to create these figures can be found in SN 15.17. To see populations overlayed onto a GF_offset_ model interpolated across the species range see Fig.

#### 3.2.2 Douglas-fir

Gradient Forest models predicting offset to future climates (RCP8.5 2050s) using WZA loci gave inconsistent results as to which set of Douglas-fir populations were projected to be most maladapted to new climates (Fig. 5). For the coastal variety, the cross-variety model and the coastal-only model of GF_offset_ each identified the same two populations from coastal BC to be the least maladapted, but rank changed considerably among the remaining populations (Fig. 5A). For the interior variety, the cross-variety and the interior-only model results conflicted as to whether the northwestern genetic group (Fig. S1) or the southeastern genetic group would be more maladapted (compare Fig. 5C and Fig. 5D), whereas these models agreed when projecting offset to the common garden (Supplemental Text S1.7; Figs. S13-S15). For the northwestern interior genetic group, results from the cross-variety and interior-only models were generally similar, except that the population identified as the least maladapted with the cross-variety model was the most maladapted from the interior-only model (Fig. 5E). For the southeastern interior genetic group, there was a negative relationship between offset predicted by the two models (Fig. 5F).

**Fig. 5.**
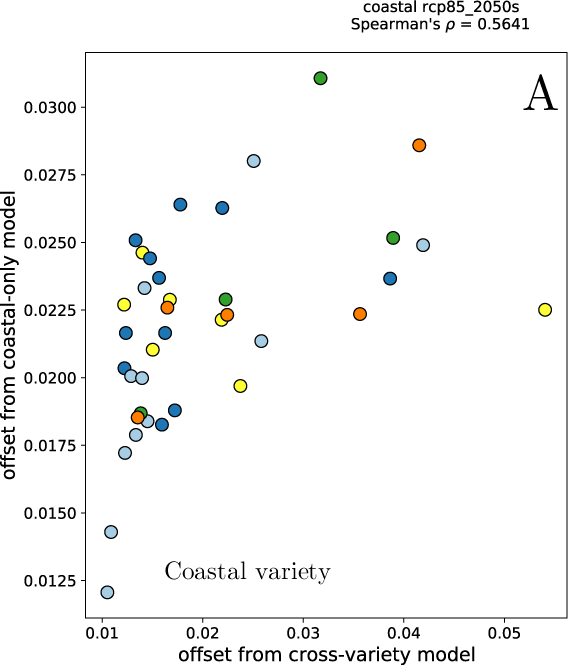
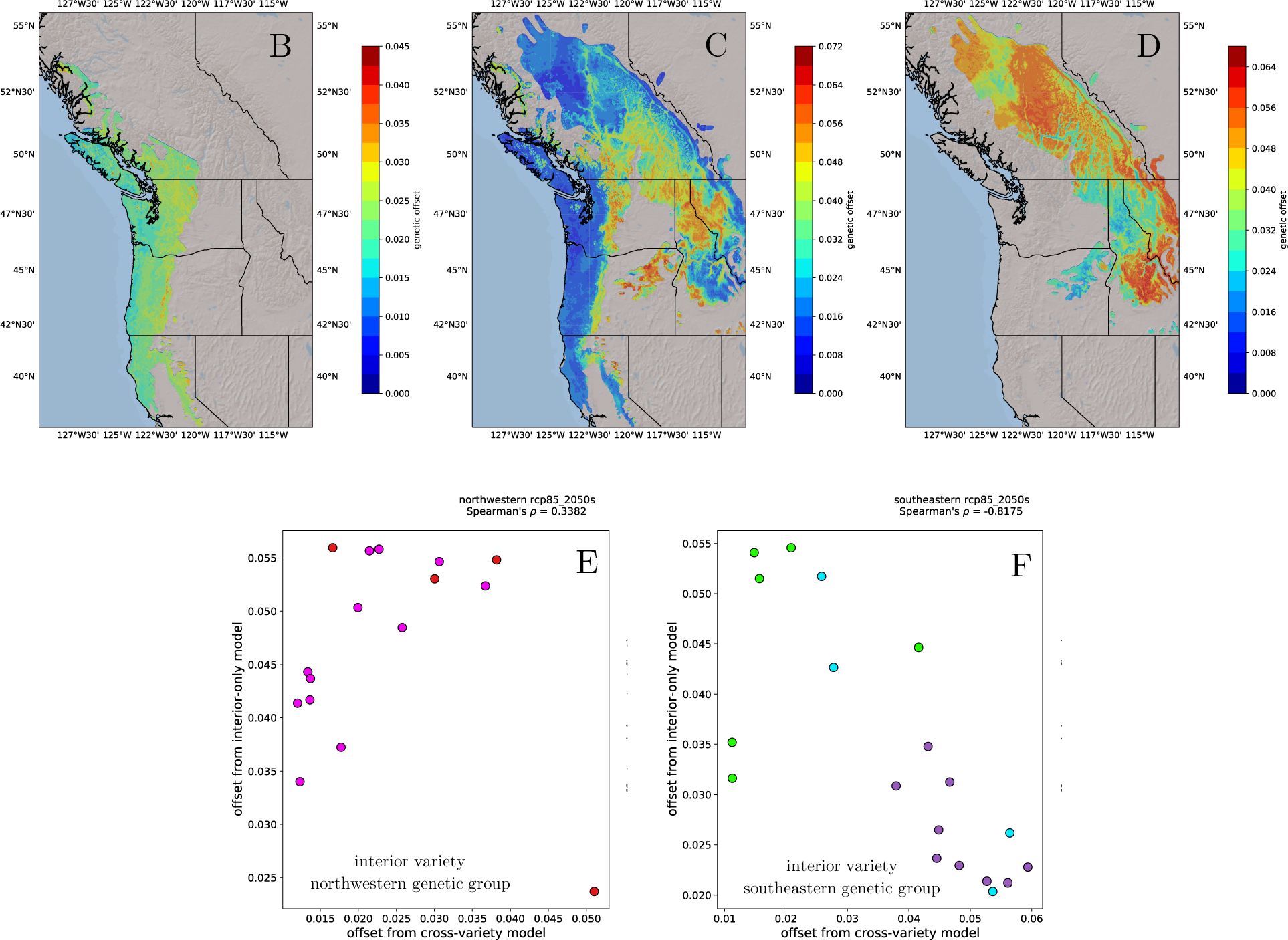
Maladaptation of Douglas-fir populations to future climate (RCP8.5 2050s) inferred from Gradient Forests (GF_offset_) is inconsistent between models trained using both varieties with those trained on a variety-specific basis. Shown are projected offsets to the range of Douglas-fir trained using WZA candidates and all populations from B) the coastal variety, C) both varieties, and D) the interior variety. For coastal Douglas-fir (A) and the two subvariety genetic groups of interior Douglas-fir (E, F), the relationship between the magnitude and rank of projected offset using the cross-variety model (pentagons, y-axes) is contrasted to those from the variety-specific model (squares, x-axes). Of note, the cross-variety model (C) and the interior-only model (D) indicate different interior variety genetic groups (populations in E or F) to be most maladapted to projected climate. Populations are colored with respect to Fig. 1. Color legend is not standardized across B, C, and D to accentuate patterns in the data (offset values are meaningless outside of the current model). Code used to create these figures can be found in SN 15.18. Analogous figures created using climate models RCP4.5 2080s, RCP4.5 2050s, and RCP8.5 2080s show similar patterns and are not shown except within SN 15.18. To see populations overlayed onto B-D, see Fig. S12.

The most maladapted interior Douglas-fir genetic group predicted from RONA was also inconsistent between the cross-variety and interior-specific models (Fig. S13B). However, RONA predictions were generally positively correlated for the interior variety and cross-variety models for the two interior genetic groups (Fig. S13C-D). Predictions from the cross-variety and coastal-only RONA models generally had positive, albeit relative weak, relationships (Fig. S13A).

To select among the models for projecting offsets to future climate for Douglas-fir, we used three criteria when comparing cross-variety and variety-specific models: 1) validation scores, 2) agreement between future offsets from GF_offset_ and RONA, and 3) agreement among RONA future offsets (Supplemental Text S1.8; Figs. S16-S21). Based on these criteria we use the cross-variety models to project maladaptation to future climate (RCP8.5 2050s; Fig. 6). For coastal Douglas-fir, many populations found along the Pacific Coast of California (orange) and Oregon (blue) had the greatest projected maladaptation (Fig. 6A). Populations from northwestern interior Douglas-fir near the Fraser River had consistently high offset ranks (red and magenta circles, Fig. 6B), whereas the remaining populations had a wide range of projected risks, and it is unclear which would be most affected by future climate. Finally, populations of southeastern interior Douglas-fir found in Idaho, Montana, and eastern Washington and Oregon had consistently greater predicted maladaptation to future climate than those found in Southeastern British Columbia (Fig. 6B).

**Fig. 6.**
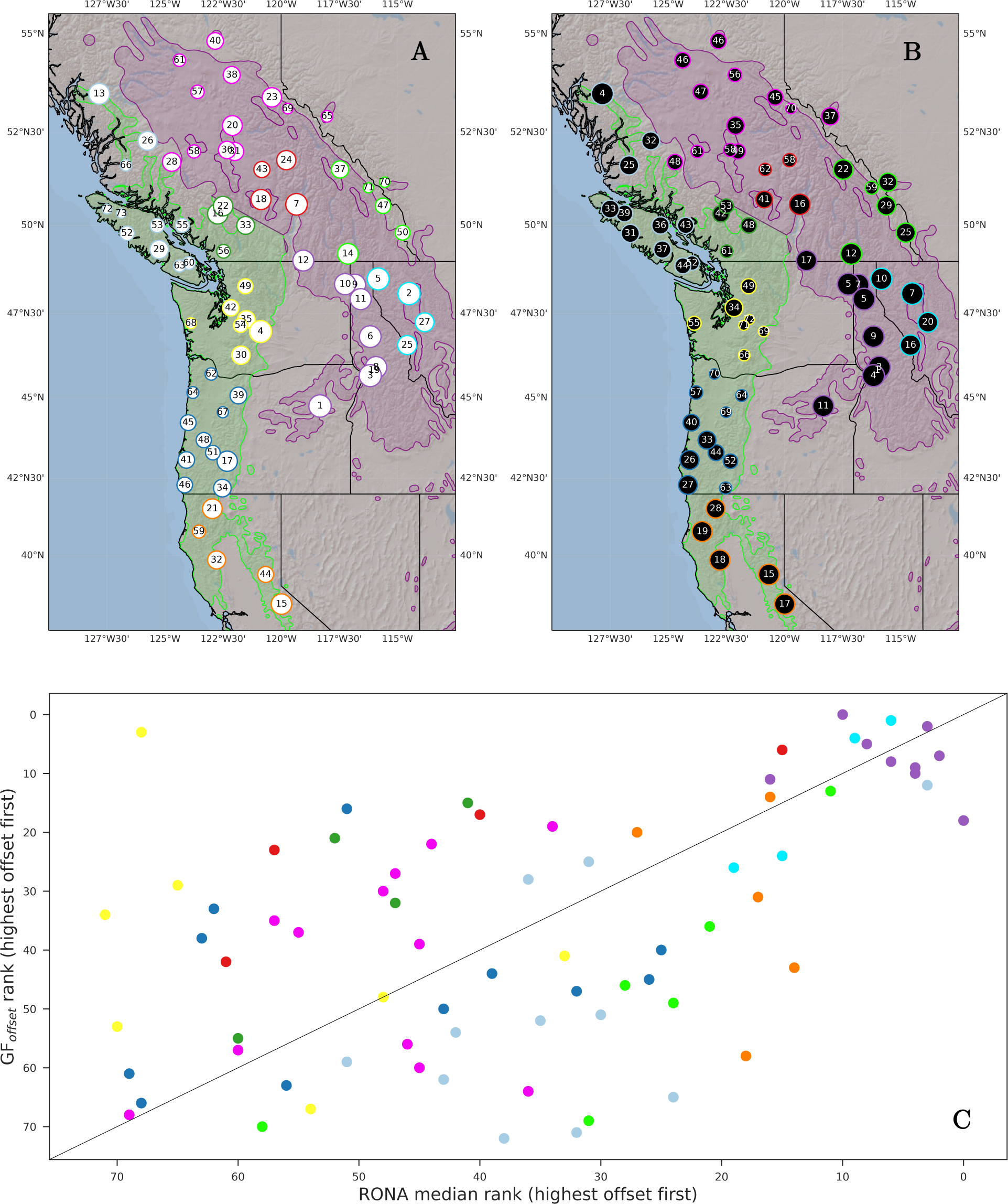
Maladaptation of interior Douglas-fir to future climate (RCP8.5 2050s) inferred from Gradient Forests (GF_offset_, A, C) and RONA (B, C) cross-variety models. Population point sizes in A and B are scaled to offset rank (lowest offset have smallest sizes). Population point sizes in B are from the median ranks across environments used to estimate RONA, which were chosen based on ranking p-values from paired *t-*tests between current and future climate. In C, a 1:1 line is given to infer relative changes in rank between methods. Rank numbers are given within circles of A and B. Populations are colored as in Fig. 1. Code to create these figures can be found in SN 15.17. To see populations overlayed onto a GF_offset_ model interpolated across the species range see Fig. S12.

## 4 Discussion

Projections of maladaptation of populations to climate change using genomic data, i.e., genomic offset estimates, have remained largely unvalidated despite the recent increase in their use. Here we use three taxa of conifers, four genomic marker sets, and common garden phenotypes from two-year Douglas-fir and 52-year jack pine individuals to demonstrate that genomic offset methods perform as well or better than the best climate or geographic distance metrics when predicting fitness-related phenotypes in transplant experiments (Q2, Fig. 2-3, Extended Data Fig. 1). We also demonstrate that candidate marker sets provide little advantage over random sets of loci (Q1). However, we find model performance at fine spatial scales was not representative of performance calculated range-wide (Q3, Figs. S9-S10). Lastly, we find that when using future climate to predict offset, the Douglas-fir populations inferred to be most maladapted depends on the model used (compare within and across Fig. 5, Fig. 6, and Figs. S13, S21). However, RONA and GF_offset_ results largely agree when projecting jack pine offset to future climates (Fig. S11). In the absence of validation data, and without further knowledge of the behavior and sensitivity of these genomic offset methods under a wider range of scenarios, it may therefore be difficult to determine whether a given set of populations can lead to reliable inferences about future maladaptation. Together, these results suggest that acting on projections of maladaptation from genomic offset methods through changes to policy and management practices should be considered only after careful scrutiny of model performance, sensitivity, and generalizability. These findings also highlight the large knowledge gap with respect to the ideal population and dataset features needed to produce reliable genomic offset models.

### Considerations for Model Construction, Validation, and Generalizability

The choice of data used to train genomic offset models, and its relationship to data used for making offset predictions, requires careful consideration and extensive exploration. A first step in model exploration is to benchmark performance with other methods that could be used to predict maladaptation. Our results mirror other studies (Fitzpatrick et al., 2021; Láruson et al., 2022) and suggest that genetic data often contains more information regarding climate adaptation than can be characterized with more readily accessible forms of data such as climate or geographic distance (Q2). This suggests that climate distance alone is unlikely to accurately estimate the extent of maladaptation of populations to future climate change.

Second, the source of inputs used to train models should be tested to understand how predictions are influenced by aspects of the source data. In our analyses, the models trained using GEA candidate loci performed no better than those from models using random loci (though there are minor exceptions for random sets used for RONA, Figs. 2-3 and Extended Data Fig. 1). This suggests it may be unnecessary to expend resources to identify adaptive genomic regions when genome- or exome-wide data exist (Q1), a finding consistent with previous evaluation of GF_offset_ (Fitzpatrick et al., 2021; Láruson et al., 2022). The similar performance among marker sets is perhaps due to the nature of our exome-targeted sequence data which targeted functionally relevant coding regions. It remains to be seen if relatively inexpensive sequencing techniques such as RAD-seq, which more often tags intergenic regions of large genomes (Parchman et al., 2018), would perform as well as the random marker sets used here. Even so, for species with strong local adaptation where isolation-by-environment drives spatial genetic structure, signals from genotyping-by-sequencing markers may contain sufficient information for accurate offset projection and may therefore be a cost-effective alternative to the exome capture data used here. Other input sources could be tested as well, such as varying the climate period used in training during model selection.

Third, the phenotypes and environments used to validate offset models should be varied to understand how performance varies with different components of fitness as well as the extent to which these predictions change with the validation environment. For example, the contrast in the performance of these offset measures across the two jack pine provenance trial sites highlights the value of using multiple sources of validation in future work, and suggests that performance may vary with validation conditions (i.e., the ‘future’ environment). Future studies will require validation to provide any degree of confidence in informing population- or site-specific management decisions. At a minimum, they will need to consider the extent to which the phenotypes and life stage used in validation are associated with total lifetime fitness (Fitzpatrick et al., 2018), as well as how the common garden environment interacts with these phenotypes. For instance, while jack pine 52-year DBH may capture elements of fitness related to growth, it may miss aspects of fitness more directly related to survival and reproduction. Size phenotypes such as DBH may also be more indicative of competitive ability in the planted common garden environment than fitness in the wild. Carefully considering the phenotype used to validate model predictions can help avoid ambiguous situations where it is unclear if poor performance is due to the choice of validation phenotype or the model itself. Varying the validation environment will also incorporate uncertainty into predictions of maladaptation to climates that may differ from those used in validation.

Fourth, the populations used to validate offset models should be relevant to the scale at which management is applied (Q3). For instance, had we chosen the cross-variety model to apply towards management recommendations in Douglas-fir, but not assessed performance at finer spatial scales, we may have concluded that the validation score from the cross-variety model was indicative of a well-performing model across all populations from both the interior and coastal varieties of Douglas-fir. However, this would have misguided prioritization of populations within these groups, as the performance of the cross-variety model decreased at the more relevant within-variety level for coastal and interior of Douglas-fir. Future sampling designs should take genetic structure into consideration and ensure that sampling is relevant to the scope of management within each genetic group. Studies should also explore the influence of highly diverged populations (e.g., those isolated from large contiguous ranges) in biasing model estimates. While it is important to consider differences between training and test data (see below), genetic and climatic differences among populations used in training should also be explored to quantify biases introduced by differentiated input data, such as with leave-one-out sensitivity analyses (Géron, 2022; Lever et al., 2016; Lotterhos et al., 2022; Rellstab et al., 2021).

Fifth, the relationship between the data used in training and prediction must be assessed to understand model generalizability and therefore the ability to make predictions on novel conditions not seen in training. Understanding model generalizability is fundamental for using predictive models, and it is well known that many mathematical models may not predict well to novel conditions relative to data used in training (Géron, 2022; Lever et al., 2016; Raschka & Mirjalili, 2019), and this applies to genomic models as well (Fraslin et al., 2022; Ma & Zhou, 2021; Rogers & Holland, 2021; Schrider & Kern, 2018; Wientjes et al., 2013). For the data used here, the transplant sites used to validate models for coastal Douglas-fir and jack pine were within the climate space of the training populations. However, the Vancouver common garden was well outside the climate space of the interior Douglas-fir populations (Fig. S23) which had the poorest performance among the three taxa assessed. While this observation could be due to weaker local adaptation in interior Douglas-fir, it may instead indicate that projections of maladaptation to future climates that differ greatly from climate data used in training may produce less robust estimates. With many marine and terrestrial environments in the mid-21^st^ century having no 20^th^ century climate analog (Lotterhos et al., 2021; Mahony et al., 2017), offset methods may be effective only for short-term predictions.

### Ignoring offset model assumptions may lead to misguided inference

Even with some promising results here, genomic offset estimates should be used with caution to guide management decisions, as there are circumstances under which these estimates may be misleading with respect to true population maladaptation even under otherwise ideal circumstances (e.g., in the presence of local adaptation). In addition to having the necessary data for accurate genomic offset predictions, not all species (or groups of populations) are ideally suited for these models. These models assume all populations are locally adapted to recent climates, that current genotype-climate relationships are due solely to local adaptation and will remain optimal in the future, and that deviations from these relationships will result in decreased fitness (Capblancq et al., 2020; Rellstab et al., 2021). Because these models assume that the change of the environment is immediate (Fitzpatrick & Keller, 2015; Láruson et al., 2022), they also ignore other dynamics that could either alleviate or exacerbate maladaptation experienced by future populations, such as gene flow (and perhaps subsequent swamping) of adaptive alleles, changes in competition or disease, or the redundancy in the genetic architecture underlying fitness and therefore the number of available routes to adaptation (Capblancq et al., 2020; Láruson et al., 2020; Rellstab et al., 2021). Because these factors could alter population trajectories between current and projected climate scenarios, offset models may be most accurate for short-term *in situ* predictions, or for predictions most relevant to near-term assisted gene flow initiatives.

### Future work is needed to identify the domain of offset applicability

There is still considerable uncertainty in the usefulness of genomic offset methods for natural populations (Capblancq et al., 2020; Láruson et al., 2022; Rellstab et al., 2021). Investigators are further limited when applying genomic offsets across taxa because the domain of applicability – i.e., the circumstances under which a method is acceptably accurate (Lotterhos et al., 2022) – remains largely undefined. For offset methods, these circumstances encompass the evolutionary history of targeted populations as well as the design of experiments used to train and validate the model itself. Even with ideal data, offset inferences will be affected by both evolutionary factors (e.g., drift, pleiotropy, and the drivers and strength of divergent selection) and experimental parameters (e.g., sampling locations, Láruson et al., 2022). The circumstances under which we should expect multiple offset methods to agree are also unclear, as they are likely to be affected to different degrees for any given set of experimental and evolutionary parameters. For example, Láruson et al. (2022) highlight how genetic drift can mislead GF_offset_ magnitude and rank estimates. This may be driving some patterns observed here, for example, the extent to which the western-most group of jack pine is inferred to be the most maladapted to future climate change (Figs. 4A, S11) or the extent to which the cross-variety model of Douglas-fir infers the southeastern groups of the interior variety to be most maladapted as well (Figs. 5C, 6B, S21.9). Because of this, we hesitate to recommend either GF_offset_ or RONA over the other, given their similar performance as well as the uncertainty of how their performance may differ in other situations. Instead, we recommend further exploration of their performance under a wide variety of scenarios, as has been noted elsewhere (Capblancq et al., 2020; Rellstab et al., 2021). A more detailed understanding of how genomic offset methods interact with complex multivariate selection, admixture, lesser degrees of (or variation in) local adaptation, and prediction to novel and strongly differentiated climates also warrant further attention.

### Concluding remarks

Ultimately, defining the domain of applicability for genomic offset methods will likely require extensive evaluation of simulated and empirical data. Until such a domain is well defined, future work estimating genomic offsets will need to thoroughly explore the results by varying input loci, climate data, populations used in training, and environments used for validation to understand how sensitive the offset estimations are to the data at hand as well as how generalizable these models are when predicting to novel data. Such exploration should follow best practices (Géron, 2022; Raschka & Mirjalili, 2019) and will require training of many dozens of models for a single dataset, which will provide ample targets for model selection and tuning. Doing so will lead to a more complete understanding of the performance of these models, and the circumstances under which they will fail.

While our validation results show promise, our future projections for Douglas-fir show ambiguous results (Figs. 5C-5D, S13B). Because of this, we do not recommend using offset estimates to strongly influence prescriptions to guide climate-adaptive management practices for individual populations until these approaches are better understood and validated. It therefore may be more prudent to work under the assumption that all populations are at some risk of maladaptation due to climate change. Even so, offset methods could guide *ex situ* conservation collections to capture genetic diversity from populations predicted to be most at risk of climate-related extirpation, e.g., for seed banks or living collections. However, because of the expectation that model performance will suffer as the environments between training and predictions diverge, we strongly caution against implementing widespread management actions based on inferences from offset models projected to climates strongly differentiated from current conditions (e.g., projections beyond several decades). While our offset projections for Douglas-fir show ambiguity in model projections, monitoring populations for climate change responses could provide evidence that support one projection over the other and provide additional validation. In practice, there may be situations where the risks of inaction may outweigh risks associated with model uncertainty, and these could be weighed accordingly, particularly for threatened or endangered species. Finally, the value of common garden experiments for evaluating risk of maladaptation should not be underestimated.

## 5 Acknowledgements

The CoAdapTree project is funded by Genome Canada (241REF; Co-Project Leaders SNA, SY and Richard Hamelin), with co-funding from Genome BC and 16 other sponsors (http://coadaptree.forestry.ubc.ca/sponsors/), including Genome Alberta, Génome Québec, the BC Ministry of Forests, and Alberta Innovates Bio Solutions. We thank Centre d’expertise et de services Génome Québec for sequencing service, University of Calgary Information Technologies for system support, as well as Dr. Pia Smets and Christine Chourmouzis for technical assistance. We also thank CoAdapTree Scientific Advisory Board members Drs. John Davis, Matias Kirst, and Graham Coop for their guidance. SY is also funded by a Natural Sciences and Engineering Research Council of Canada (NSERC) Discovery Grant (RGPIN/03950-2017) and Alberta Innovates (20150252), and SA is funded by an NSERC Discovery Grant (RGPIN-2020-05136). The funding bodies did not have any role in the design of the study, collection, analysis, or interpretation of data in writing the manuscript. Finally, we thank Katie Lotterhos and reviewers Thibaut Capblancq, Christian Rellstab, and an anonymous reviewer for very helpful comments to improve our manuscript.

## 7 Data Availability

We reference the analysis code in the text of our documents by designating Supplemental Notebooks (SN) using a directory numbering system from our servers (as opposed to the order listed in the manuscript). For example, for Notebook 15 in Directory 3, we will refer to SN 03.15; for Notebook 1 in Subfolder 2 of Directory 5, we will refer to SN 05.02.01. These notebooks not only contain the analysis code, but also contain code output and display attributes of the data objects being analyzed. Each of these directories are archived on Zenodo.org and include a citation below, which will also link to the GitHub repository. Notebooks are best viewed within a local jupyter or jupyter lab session, but can also be viewed at nbviewer.jupyter.org using the web link to the notebook within the archived README (also available on GitHub). Analyses were carried out primarily using python v3.8.5 and R v3.5.1. Exact package and code versions are available at the top of each notebook. Anaconda environments used to carry out analysis can be recreated using the .yml files found in the archives.

Archives of Supplemental Notebooks:

SN 15 : Lind, B.M. 2023. GitHub.com/brandonlind/offset_validation: Revision 1 release. Zenodo. https://doi.org/10.5281/zenodo.7641225

SN 02 : Lind, B.M. 2023. GitHub.com/brandonlind/ douglas_fir_natural_populations: Offset Revision 1 (v1.0.0). Zenodo. https://doi.org/10.5281/zenodo.8018894

SN 07 : Lind, B.M. 2023. GitHub.com/brandonlind/jack_pine_natural_populations: Offset Revision 1 (v1.0.0). Zenodo. https://doi.org/10.5281/zenodo.8018892

Raw sequence data will be deposited on the Sequence Read Archive of the National Center for Biotechnology Information (NCBI SRA). All remaining data necessary for the replication of our work will be archived on DataDryad.org.

## 8 Author Contributions

SA and SY obtained funding. BL and SA conceived the offset validation study. RC-R and NI provided phenotypes used in validation for Douglas-fir and jack pine, respectively. PS carried out gene annotation. TB implemented WZA, which was designed by TB, MW, and SY. BL processed raw genetic data, called SNPs, and carried out single-locus GEA for each species. BL carried out implementation of genomic offset training, validation, and projection. BL wrote the manuscript, with editing and feedback from all authors. BL created figures, and curated coding records for archiving. All authors contributed to improvements of the study design and the conceptualization of results.

## 9 Extended Data

**Extended Data Table 1.** Locus counts used in training of Gradient Forests (GF_offset_) and the Risk Of Non-Adaptedness (RONA) for each set of populations used: jack pine, across both varieties of Douglas-fir (cross-variety), coastal Douglas-fir, and interior Douglas-fir. Not shown are redundant counts of random marker sets with the same sample size as the BayPass and WZA sets used in GF_offset_. Marker sets used for RONA are subsets of those used in GF_offset_ that had significant linear models with at least one environment. Subscript letters b and w refer to the original candidate sets (BayPass, and WZA, respectively) used to determine sample sizes for random marker sets used in GF_offset_ which were then subset to form the counts shown for RONA. Code used to create this table can be found in SN 15.15.

|  | Gradient Forests |  | RONA |  |  |  |
| --- | --- | --- | --- | --- | --- | --- |
|  | BayPass | WZA | BayPass | random<sub>b</sub> | WZA | random<sub>w</sub> |
| Jack pine | 22635 | 8564 | 22570 | 11383 | 8563 | 4281 |
| Cross-variety | 25219 | 4810 | 24687 | 22857 | 4756 | 4337 |
| Coastal | 17516 | 3770 | 17433 | 9684 | 3766 | 2050 |
| Interior | 12938 | 1787 | 12262 | 5973 | 1787 | 873 |

**Extended Data Fig. 1.**
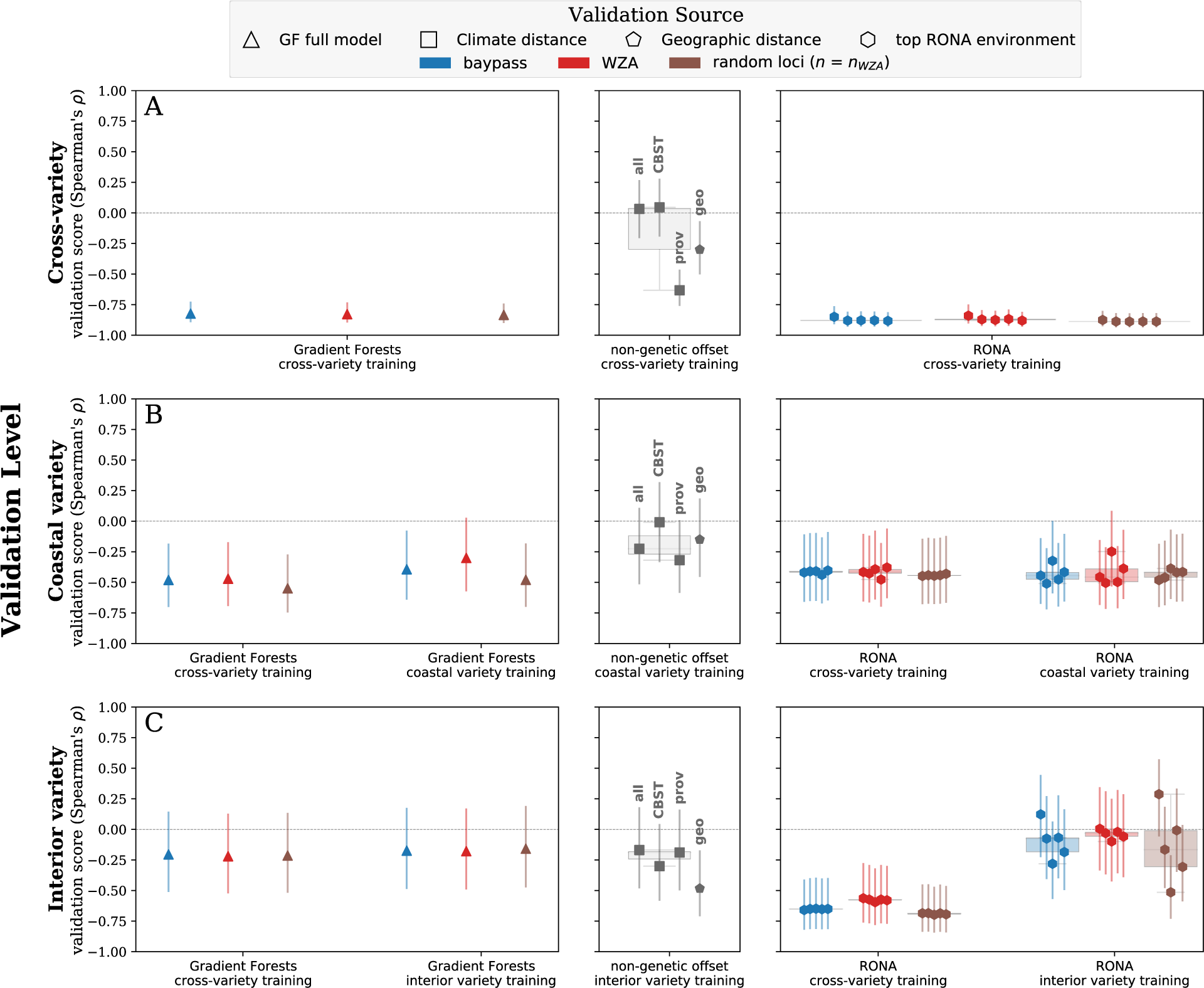
Offset validation from two-year Douglas-fir shoot biomass phenotypes at the Vancouver common garden (see Fig. 1A) using Gradient Forests (GF_offset_), the Risk of Non-Adaptedness (RONA), and climate and geographic distances. We used genetic hierarchy (rows) to assess accuracy inference from trained models (x-axis groups) using populations (rows) across both varieties of Douglas-fir (A), at the variety level for the coastal (B) and interior varieties of Douglas-fir (C) to determine if greater numbers of training populations improve finer-scale predictions of offset. Triangles indicate performance of GF_offset_ models trained and validated using all available populations. RONA background boxplots illustrate the range of RONA validation scores given for the top five climatic variables (hexagons) that differed significantly between source population and the common garden (see Table S1). Climate distances (squares) were calculated using 1) all climate variables, or 2) those variables used for climate-based seed transfer (CBST) in British Columbia, or 3) those explaining significant variation in provenance trials. Vertical bars indicate standard error estimated using a Fisher transformation (see Supplemental Text S1.3). Loci used in RONA calculations are a subset of those used in Gradient Forests that had significant linear models with the environment, see Table 1 for locus counts. See Fig. 2 for similar validation using height increment. See Fig. S9 for all locus groups. Boxplot whiskers extend up to 1.5x the interquartile range. Code to create these figures can be found in SN 15.14.

**Extended Data Fig. 2.**
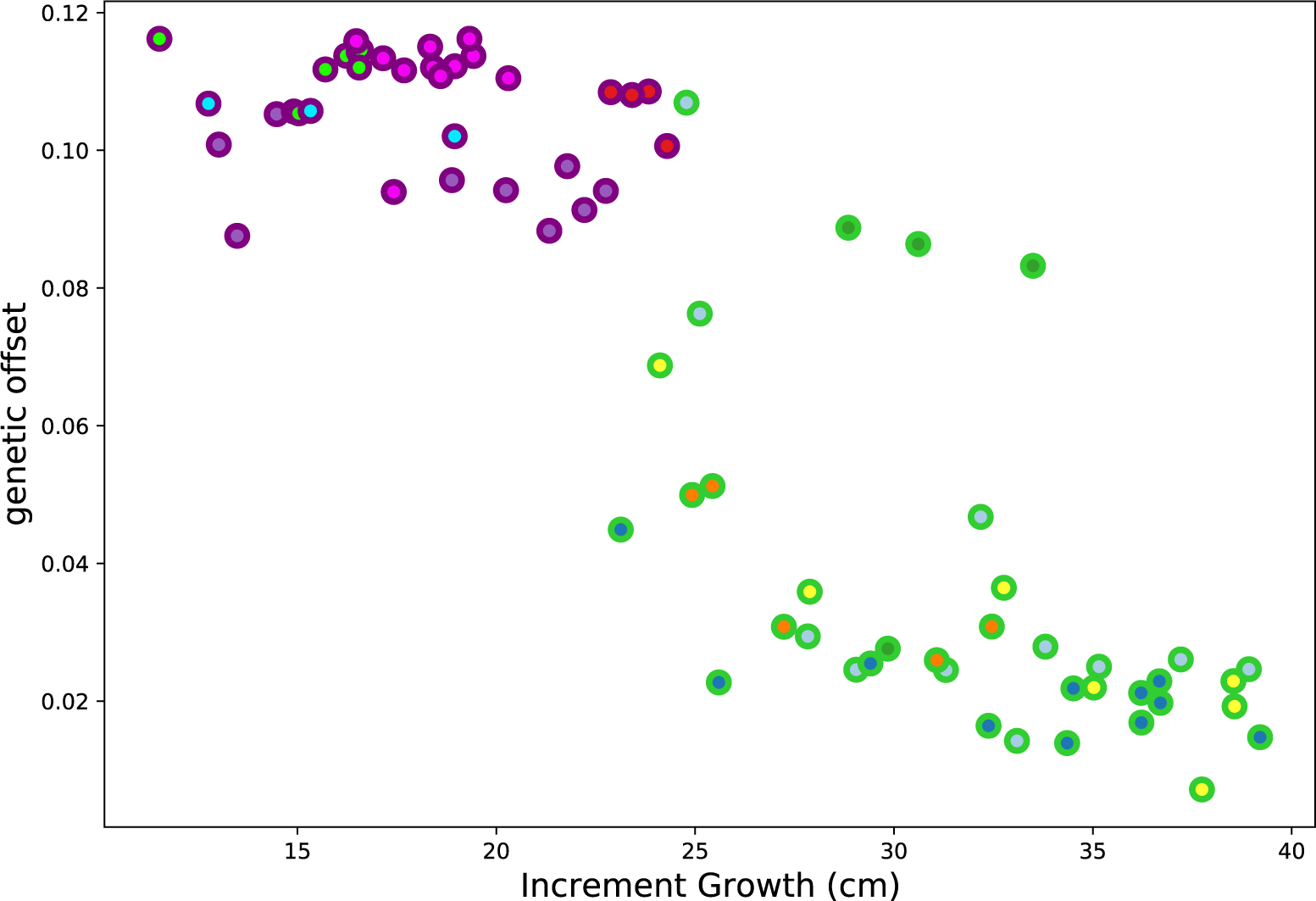
Genetic structure drives high validation scores in the Douglas-fir cross-variety models of Gradient Forest (GF_offset_). Shown is the relationship between increment growth of coastal and interior varieties of Douglas-fir and offset values from the cross-variety model of GF_offset_ (trained using WZA candidates and all populations). Edges are colored using the shade of each variety’s geographic range in Fig. 1 (lime: coastal Douglas-fir, and purple: interior Douglas-fir), and the interior of each point is colored with respect to the genetic groups from Fig. 1. Code used to create this figure can be found in SN 15.22.

**Extended Data Fig. 3.**
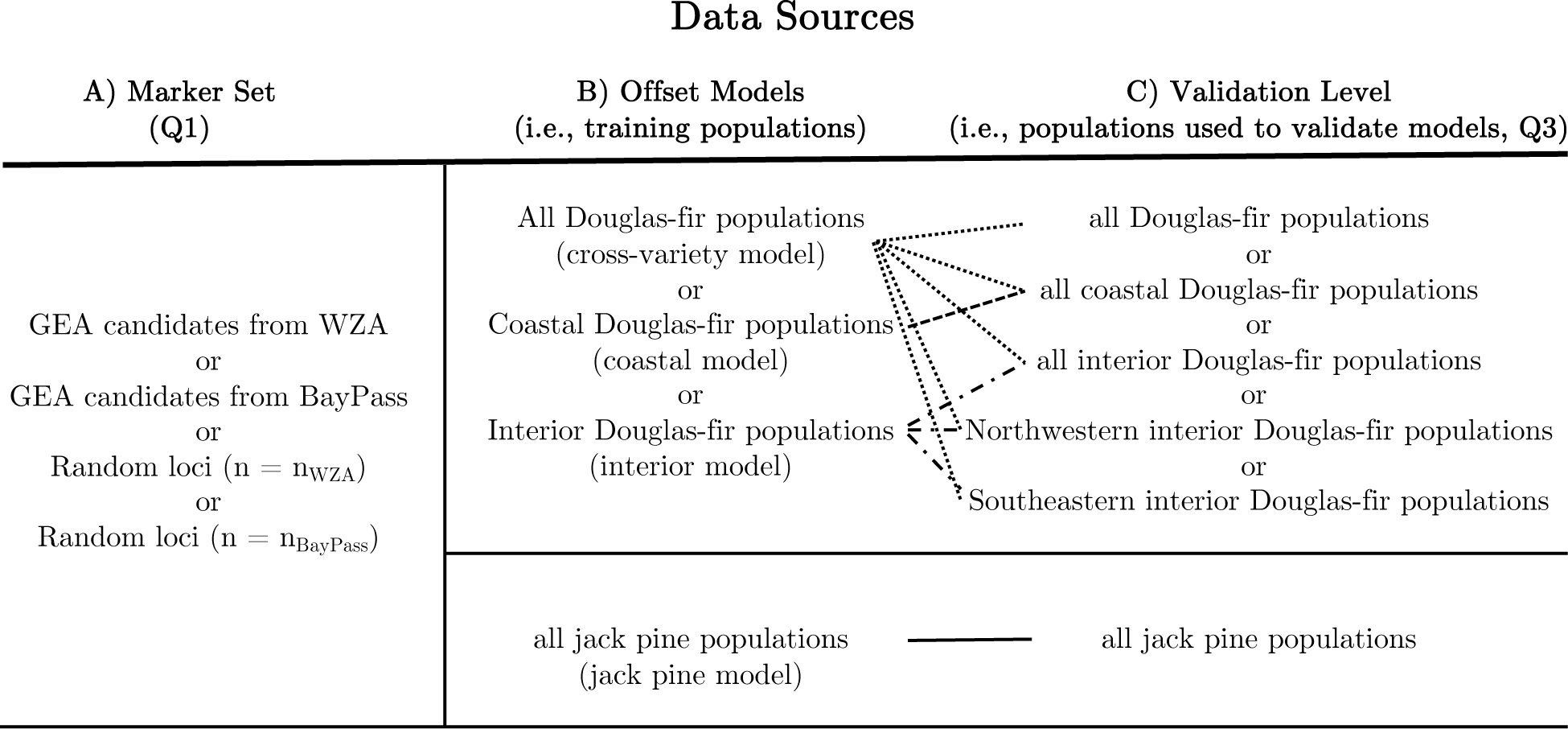
Relationship between data used to train and validate genomic offset models. Marker sets were varied to understand impact of marker source (A, Q1). Populations from Douglas-fir and jack pine were used to create four sets of training populations to train offset models (B). Offset models were validated using either all populations used in training or subsets of these populations (C, Q3); lines connecting (B) to (C) indicate which population subsets in (C) were used to validate models in (B). Not shown are population sets used to understand model generalizability (Q4). The *coastal and **interior models are sometimes referred to as coastal-only or interior-only models for readability.

