## supplement for "How useful is genomic data for predicting maladaptation to future climate?"

30 August 2023

**Running Title**: *Validating genomic offsets*

**Keywords**: climate vulnerability, climate change, machine learning, genomic offset

[*]**Corresponding Author**

Brandon Lind

2424 Main Mall

3027 Forest Science Centre

University of British Columbia

Vancouver, BC, Canada

### 1 | Supplemental Text

#### 1.2 | Genetic grouping

We identified genetic groups as informed by PCA *k*-means clustering (Fig. S1-S2) to color populations for visualization purposes. To choose the optimal number of genetic groups we used a combination of the mean silhouette coefficient scores from first three axes of PCA fit using loci with no missing data and the second derivative of the line fit (hereafter, Line 1) using the Number of Clusters and the Within-Cluster Sum of Squares (SN 07.04.01, SN 02.12.01, SN 02.12.03). The within-cluster sum of squares used here is the unweighted sum of squared distances of samples to their closest cluster center for a given value of $k$. The silhouette coefficient is calculated on a per-sample basis using the mean intra-cluster distance and the mean nearest-cluster distance; the best value is 1 and the worst value is -1. We calculated mean silhouette coefficient scores for $k$=2-20, each time recording the within-cluster sum of squares, and chose the optimal number of $k$ when the mean silhouette coefficient first reached a local maximum (for any $k > 2$) and Line 1 appeared to have a second derivative close to zero. For Douglas-fir varieties, this generally resulted in less than three optimal clusters; for the interior variety this was the northwestern and southeastern genetic groups, and for the coastal variety this was the central range and range-edge populations. So we used information from the PCA, *k*-means clustering, and geography (see subfolder 02.03) to increase the number of groups, in line with groupings found when using larger values of *k* during *k*-means clustering of PCA data. PCAs colored with according to the *k*-means clustering (i.e., colors in Fig. 1) can be seen in Fig. S1 for Douglas-fir and Fig. S2 for jack pine.

##### 1.3 | Consistency of Environmental Variable Importance Rank

The environmental variables driving local adaption are often unknown *a priori*, and landscape ecologists must often choose among highly correlated environmental variables when analyzing data. Random forest models, like those implemented within Gradient Forests, are well suited to identify features important for predictive models, even when presented with correlated features. In the context of GF, these features are environmental variables. We assessed the feature importances output by GF using both accuracy importance and weighted accuracy importance and visualized the consistency of these importance ranks from each dataset using slope graphs. If the various marker sets and populations used to train GF models have little impact with respect to climate feature importance, ranks should be stable across these outcomes and therefore these slope graphs should contain relatively little noise (i.e., no large or numerous changes in rank), at least for the top-ranked climates which may contain considerably larger importance values than lower ranked climate features. If these figures are noisy throughout all ranks, this tells us that the input data impacts model outcomes with respect to inferred climate importance.

Across the Douglas-fir models trained using BayPass or WZA candidate s, top environmental ranks from weighted importance values were most similar across runs between the cross-variety and interior-only models, where TD, EMT, and MCMT were ranked in the top four environments across runs (Fig. S7A-C). For the coastal variety models trained using candidate sets, Eref, CMD, EXT, MSP, and SHM were consistently ranked as the top five variables across runs (Fig. S7B). For jack pine, the environmental ranks were the same except the rank was switched between DD5 and EMT (Fig. S7D).

For all Douglas-fir models trained using candidate s and all populations (blue circles and diamonds, Fig. S7), the top six environments were consistently ranked between BayPass and WZA candidate sets. Together with the environments consistently ranked across runs, these data suggest that the interaction of summer precipitation and temperature play an important role in structuring sampled genetic variation in the coastal variety, while cold temperatures predominate as important factors structuring sampled genetic variation within the interior variety. Similar to Douglas-fir, models trained using all jack pine populations and loci from BayPass and WZA candidate s produced consistent rank and indicate both cold temperatures and precipitation drive patterns within our genetic data for jack pine (Fig. S7). For both species, the ranks from weighted importance were in general agreement with those from accuracy importance (Fig. S5).

Except for the GF model trained using the coastal variety, GF models generally ranked the majority of CBST environmental variables within the top 50% of ranks (blue and red environmental labels in Fig. S7, Fig. S5). However, environmental variables explaining significant variation in Douglas-fir provenance trials were generally in the bottom 50% of ranks, except for MAT in the interior-only models (brown and red environmental labels in Fig. S7A-C, panels A-D in Fig. S5). For jack pine, the provenance trial environments (except for SHM) generally ranked in the top 50%, where environments related to both CBST and provenance trials generally predominated top ranks (brown, red, and blue labels in Fig. S7D, panels E-F in Fig. S5). In general, the magnitudes of weighted importance indicated that between 1 to 5 climate variables could be inferred to have the greatest importance in these models (Fig. S6; analogous histograms to Fig. S6 for accuracy importance show similar patterns and are not shown, except in SN 15.13). Given the relatively consistent rank across these top-ranked climate features, our analyses indicate that both marker and population input has relatively less impact on climate importance inference than it does when predicting genetic offset.

Our analyses suggest that inference related to climate importance is relatively less sensitive than offset predicted from models of GF, though whether the most important climates ranks indicate the climates most relative to adaptation, as opposed to just patterns from the data, remains to be seen and warrants further attention.

#### 1.4 | Common garden phenotypic information for Douglas-fir and jack pine

##### *1.4.1 | jack pine*

The *Pinus banksiana* genetics program was initiated by Petawawa Experimental Forest Research in 1950. Between 1955-1961, seeds were collected by Canadian Forest Service (CFS) and United States (US) collaborators from 99 *P. banksiana* provenances throughout the species' natural range (Yeatman 1974). In 1962, seeds were sown and seedlings were then grown and evaluated at the Petewawa nursery. Three to four years later, twelve field tests, with varying numbers of provenances, were established in Canada (Ontario, Quebec, and New Brunswick) and the US to assess the pattern of adaptive genetic variation (Yeatman 1974).

In September 2018, a census was conducted in two Quebec field tests. Tree status (e.g. alive or dead) was noted along with tree height (m) and diameter at breast height (cm) for each of the trees still present. Provenance numbers 73 and 74 were both assigned to Marl Lake with the same LAT and LONG coordinates, had the same mortality levels, and similar means for height and diameter at breast heigh (DBH) – we therefore averaged these populations for height and DBH and assigned these values in the data to Provenance 73 and removed Provenance 74.

Since we were using a trial planted long ago, we matched provenance ID (LAT and LONG) used for phenotyping with that from genomic populations used for offset training by ensuring each provenance ID was < 1km from only one other population ID from our pool-seq data (SN 15.06; yellow- and black-edged circles, Fig. 1B); the population-provenance pairs that were kept all had a distance of 0.0 km, except for one pair that had a distance of 0.048 km.

*1.4.2 | Douglas-fir*

The Douglas-fir common garden experiment was established in outdoor nursery raised beds at the University of British Columbia campus in March 2018. One-year-old seedlings of 73 natural populations (38 var. *menziesii* and 35 var. *glauca*) spanning most of the natural range of the species (see Fig. 1A) were randomized into 11 blocks with an unbalanced block design. Each block contained 240 seedlings surrounded by a row of unmeasured edge seedlings. We used a spacing of 8 x 8 cm between seedlings and between one and 13 seedlings per provenance in each block. The total number of seedlings per provenance in the experiment varied between 11 and 95. All individuals in the experiment were assessed for initial height in 2018 (before bud flush of the second growing season) and final height in 2019 (after bud set of the third growing season), and final total shoot dry biomass.

Best linear unbiased estimates (BLUEs) of the two-year height increments were obtained for provenances before testing for associations with the genetic offsets. Height increments were first log transformed to meet the assumptions of normality of residuals and homoscedasticity of variances in the models. The following mixed effect model implemented in ASReml-R 4.0 (Butler et al. 2007) was used:

$$Y_{ij} = \mu + \beta x_1 + \alpha x_2 + \epsilon_{ij}$$

where $Y_{ij}$ is the phenotype height increment corresponding to individuals from provenance *i* and block *j*; $\mu$ is the phenotype global mean across all individuals within the experiment (fixed intercept), $\beta$ is the coefficient for the fixed effect of provenance ($x_1$) and $\alpha$ is the random effect of blocks ($x_2$) and $\epsilon_{ij}$ is the error term for $Y_{ij}$.

##### 1.5 | Estimating error in Spearman's $\rho$ estimates

We used Fishers Transformation to estimate the lower and upper confidence intervals of Spearman's $\rho$ estimates used in validation via the following equations:

$$F(\rho) = artanh\,(\rho)$$

$$SE = \frac{1}{\sqrt{N-3}}$$

$$lower = tanh\,(F(\rho) - (1.96 * SE))$$

$$upper = tanh\,tanh\,(F(\rho) + (1.96 * SE))$$

Where $F(\rho)$ is Fisher's transformation of Spearman's $\rho$, and SE is the standard error standardized using the number of populations used in the correlation estimate (N). Some cross-validation scores for Douglas-fir were equal to $\rho = -1.0$ (Supplemental Figs. S9D-E and S10D-E) and in these cases we set $\rho = -0.9999999999999999$ to avoid domain errors when calculating $F(\rho)$.

##### 1.6 | Exploring fine-scale validation of models trained across larger spatial scales

We continued exploring the effects of fine-scale validation (see Supplemental Note S1.10) by subsetting the interior variety into its two distinct genetic groups (the northwestern group and the southeastern group, see Supplemental Fig. S1) to use for validation from models trained across both varieties or those trained using all interior populations. While the GF models using all interior populations in validation (triangles, Fig. 3C) were in some cases outperformed by a non-genetic offset measure, validation scores using either of the two interior genetic groups often exceeded these same climate and geographic distance measures (Fig. S9D-E). Furthermore, these sub-interior variety scores (Fig. S9D-E) were often stronger than scores obtained at the interior level (Fig. 3C), and even exceed those from the coastal variety (Fig. 3B). However, cross-variety and interior-only models showed similar performance (Fig. S9C-E). Boxplots in Fig. S9D-E span a wide range, due in part to the small sample sizes of hold-out validation assignments from stratified sampling assigned from cross-variety and interior-only models (colors, Fig. 1A). Similar patterns were found for shoot biomass (Fig S10).

##### 1.7 | Inconsistency between projected maladaptation between common garden and future climates for interior Douglas-fir

The relative rank of projected maladaptation between the two genetic groups within interior Douglas-fir was inconsistent between that predicted for the Vancouver common garden and for future climate. For projections of maladaptation to future climate, the cross-variety and interior-only models of both GF (Fig. 5C-D) and RONA (Supplemental Fig. S13B) disagreed as to whether the northwestern genetic group was more maladapted to RCP 8.5 2050s than the southeastern genetic group. However, for projections to the common garden (i.e., models depicted in Fig. 3 and Extended Data Fig. 1) the cross-variety model and interior-only model of GF were in agreement (Fig. S14) and were in disagreement for RONA (Fig. S15). It is unclear why such a 'flip-flop' would occur between cross-variety and interior-only GF models for future climate but not for this model when GF projected to the Vancouver common garden. In each case, we used the same GF model object from R and then input either the Vancouver climate or the future climate (SN 15.07). The difference between these input climates was of course the values used, but in the case of the Vancouver climate input, all of the populations had a single value for a given climate variable. Other than this, it is unclear what could be causing these differences. It may be because of the differences between current and future (common garden or RCP) climates, where the future climates are more differentiated than that of the common garden from current values. Or perhaps ancestry violating model assumptions (the northwestern interior genetic group shares secondary contact with the coastal variety). Future simulation work could explore the effects of ancestry or climate dissimilarity from current values to determine if such patterns would impact projected offset rank, but this task is beyond the ability of the sparse pool-seq data available here.

#### 1.8 | Choosing future offset models to predict the most maladaptive populations

Based on the similar level of validation performance among marker sets of either candidate or random loci for both jack pine and Douglas-fir, we explored offset to future climate using WZA candidate loci. Sensitivity of this model to population input was similar to other locus sets (SN 15.21).

Next, for Douglas-fir, we needed to decide between the cross-variety or variety-specific models (i.e., which input populations to use) to project offset to future climate change. We used four criteria to choose among models by using insight from model validation and projection: 1) validation scores, 2) the strength of the relationship between offset predicted from GF with that of RONA, 3) the strength of the relationships among RONA estimates from the top environments differentiated between current and future scenarios.

1) While the cross-variety and variety-specific models of GF had similar performance when validated at the variety level, the cross-variety model of RONA had stronger validation scores for the interior variety populations than did the interior-only model (Fig. 3, Extended Data Fig. 1). We therefore use cross-variety models to project maladaptation to future climate (RCP 8.5 2050s; Fig. 6).

3) For both the coastal and interior variety, and the genetic groups from within the interior variety, the offset predicted from the GF cross-variety models had a stronger

relationship with that predicted from RONA cross-variety models than the comparison between GF and RONA variety-specific models (Fig. S21).

4) Furthermore, models for interior Douglas-fir had stronger relationships among RONA estimates from cross-variety models than from interior-only models (Supplemental Fig. S21 iii-viii).

It is worth noting that the cross-variety model ($n_{\text{populations}} = 73$) may be less sensitive than the interior-only model ($n_{\text{populations}} = 35$) due to the relatively greater number of input populations. However, given the less sensitive behavior of coastal-only models ($n_{\text{populations}} = 38$) relative to interior-only models, where input population numbers are more similar between varieties relative to either variety and the cross-variety models, the population numbers input to interior variety models may not be completely driving sensitivity patterns seen here.

#### 1.9 | Implementation of BayPass

We used structure correction within BayPass to isolate loci putatively underlying adaptation to the environment. We used a subset of the loci used in all other analyses to estimate the covariance matrix used for structure correction. Specifically, we first randomly selected one locus per genomic reference contig longer than 1kb that had no missing data across populations and a minimum allele depth (per locus per population) of 20 and maximum allele depth of 1000 (note that for the full datasets, more relaxed thresholds for depth – $\geq 8$ – were used, see Methods). We attempted to further filter loci to those without missing data, but this ultimately skewed minor allele frequency distributions compared to the distributions of the full dataset, so we maintained missing data $\leq 25\%$. We then performed linkage disequilibrium pruning by removing one locus from any pair that together had a squared Pearson $r > 0.3578$ (coastal Douglas-fir), $r > 0.3421$ (interior Douglas-fir), and $r > 0.3757$ (jack pine) – these correspond to, respectively, the $99.5^{th}$, $99^{th}$, and $99.9^{th}$ percentiles of each distribution of pairwise r values among loci. Attempts to prune loci below these thresholds caused non-convergence of replicate estimates of the covariance matrix (not shown). At each stage of filtering, the minor allele frequency spectra were compared between the full dataset and the filtered SNPs to ensure no major biases were introduced (see Supplemental Notebooks, cited below).

We used five independent estimates of the covariance matrix to ensure convergence (all 5-choose-2 comparisons were highly correlated, Pearson's $r > 0.972$ for coastal Douglas-fir, $r > 0.965$ for interior Douglas-fir, and $r > 0.907$ for jack pine). Because each matrix was highly correlated, we took the average matrix as input to BayPass GEA – this average matrix was also highly correlated with each of the five independent estimates (Pearson's $r > 0.989$ for coastal Douglas-fir, $r > 0.991$ for interior Douglas-fir, $r > 0.964$ for jack pine). We then used this matrix in BayPass to control for structure when identifying GEA candidates. We used default settings for baypass except for seed and the following: `-d0yij 8 -pilotlength 1000 -nval 50000 -npilot 40`.

The filtering, MAF comparison between filtered and original loci at each stage, matrix estimation (including visualization of correlations among replicate matrices), and execution of GEA described above was implemented in SN 02.02.01_01 for coastal Douglas-fir, SN 02.02.01_02 for interior Douglas-fir, and 07.02.01 for jack pine. The information was gathered and combined into a single dataframe in SN 02.02.02_01, 0.2.02.02_02, 07.02.02, respectively. GEA candidates at the variety level for Douglas-fir and species level for jack pine were identified as those with mean Bayes factor ≥ 15 across the five independent chains (SN. 15.04).

#### 1.10 | Structure in the data can misguide inference related to performance

Our validation statistic, Spearman's rho, was used to infer model performance, with large magnitudes of rho indicating greater model performance (e.g., large negative values when correlating offset and performance, or large positive values when correlating offset and mortality). However, when structure exists within the data (such as genetic structure), the statistic can be misleading with respect to the scales that are of management relevance.

SN 15.23 gives a toy example of how structure could lead to large magnitudes of Spearman's rho without any explanatory ability of the model at fine spatial scales. In the example, I create two datasets – each dataset has 100 'x' values and 100 'y' values – by randomly choosing a number between 0 and 1 from a uniform distribution for both 'x' and 'y'. For one of the datasets, I add 1.0 to each value of 'x' and 'y' to create "structure" in the data (see Figure S1.10 below).

The correlation statistic across these two meaningless datasets is rho=0.74, which would under normal circumstances be considered to be indicative of a well-performing model. In the context of a validation trial, a researcher that does not explore the data could conclude that this statistic is sufficient evidence that the model performs well. However, there is no explanatory value within the two groups – within each group rho $< 0.064$.

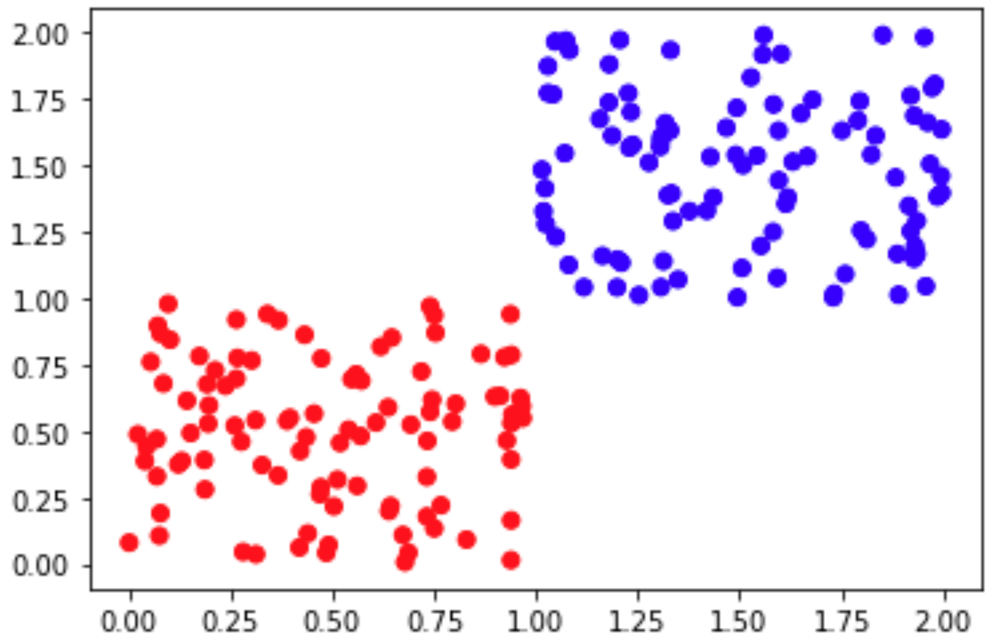


Figure S1.10 A toy example figure displaying two datasets with structure. Compare to Extended Data Fig. 2 in the main text.

### 2 | Supplemental Tables

| Environment | p-value | Common Garden | Population set |
|---|---|---|---|
| EXT | 1.084316e-27 | Ste-Christine-d'Auvergne | jack pine |
| MSP | 9.095524e-16 | Ste-Christine-d'Auvergne | jack pine |
| MAP | 5.559867e-12 | Ste-Christine-d'Auvergne | jack pine |
| SHM | 5.607879e-10 | Ste-Christine-d'Auvergne | jack pine |
| AHM | 2.107018e-08 | Ste-Christine-d'Auvergne | jack pine |
| EXT | 1.229238e-25 | Fontbrune | jack pine |
| MSP | 1.640940e-07 | Fontbrune | jack pine |
| SHM | 3.166373e-06 | Fontbrune | jack pine |
| AHM | 2.229745e-05 | Fontbrune | jack pine |
| MAP | 2.495356e-05 | Fontbrune | jack pine |
| bFFP | 3.134143e-44 | Vancouver | cross-variety |
| FFP | 6.294516e-37 | Vancouver | cross-variety |
| NFFD | 2.498133e-27 | Vancouver | cross-variety |
| eFFP | 9.501555e-27 | Vancouver | cross-variety |
| DD5 | 4.645407e-26 | Vancouver | cross-variety |
| bFFP | 1.312699e-21 | Vancouver | coastal variety |
| FFP | 1.996474e-18 | Vancouver | coastal variety |
| eFFP | 2.866360e-12 | Vancouver | coastal variety |
| NFFD | 3.582954e-12 | Vancouver | coastal variety |
| EMT | 6.254842e-12 | Vancouver | coastal variety |
| bFFP | 3.205920e-32 | Vancouver | interior variety |
| FFP | 1.292912e-31 | Vancouver | interior variety |
| eFFP | 6.564966e-30 | Vancouver | interior variety |
| NFFD | 8.936509e-30 | Vancouver | interior variety |
| TD | 1.051689e-29 | Vancouver | interior variety |

**Table S1** Top five environments that differed between source populations and common gardens ranked by t-test p-values. These ranks were used to decide which locus-environment relationships to use when calculating the range of RONA (Figs. 2-4 of the main text). Data used to create this table is in SN 15.09 cell 67 .

**Annual variables:**
MAT - mean annual temperature (°C)
MWMT – mean warmest month temperature (°C)
MCMT – mean coldest month temperature (°C)
TD – temperature difference between MWMT and MCMT, or continentality (°C)
MAP – mean annual precipitation (mm)
MSP – May to September precipitation (mm)
AHM – annual heat-moisture index (MAT+10)/(MAP/1000)
SHM – summer heat-moisture index MWMT/(MSP/1000)

**Derived annual variables:**
DD0 – degree-days below 0°C
DD5 – degree-days above 5°C
NFFD – number of frost-free days
FFP – frost-free period
bFFP -the day of the year on which FFP begins
eFFP – the day of the year on which FFP ends
PAS – precipitation as snow (mm) between August in previous year and July in current year
EMT – extreme minimum temperature over 30 years
EXT – extreme maximum temperature over 30 years
Eref – Hargreaves reference evaporation (mm)
CMD – Hargreaves climatic moisture deficit (mm)

**Table S2**. Climate variables used for offset calculations.

#### 3 | Supplemental Figures

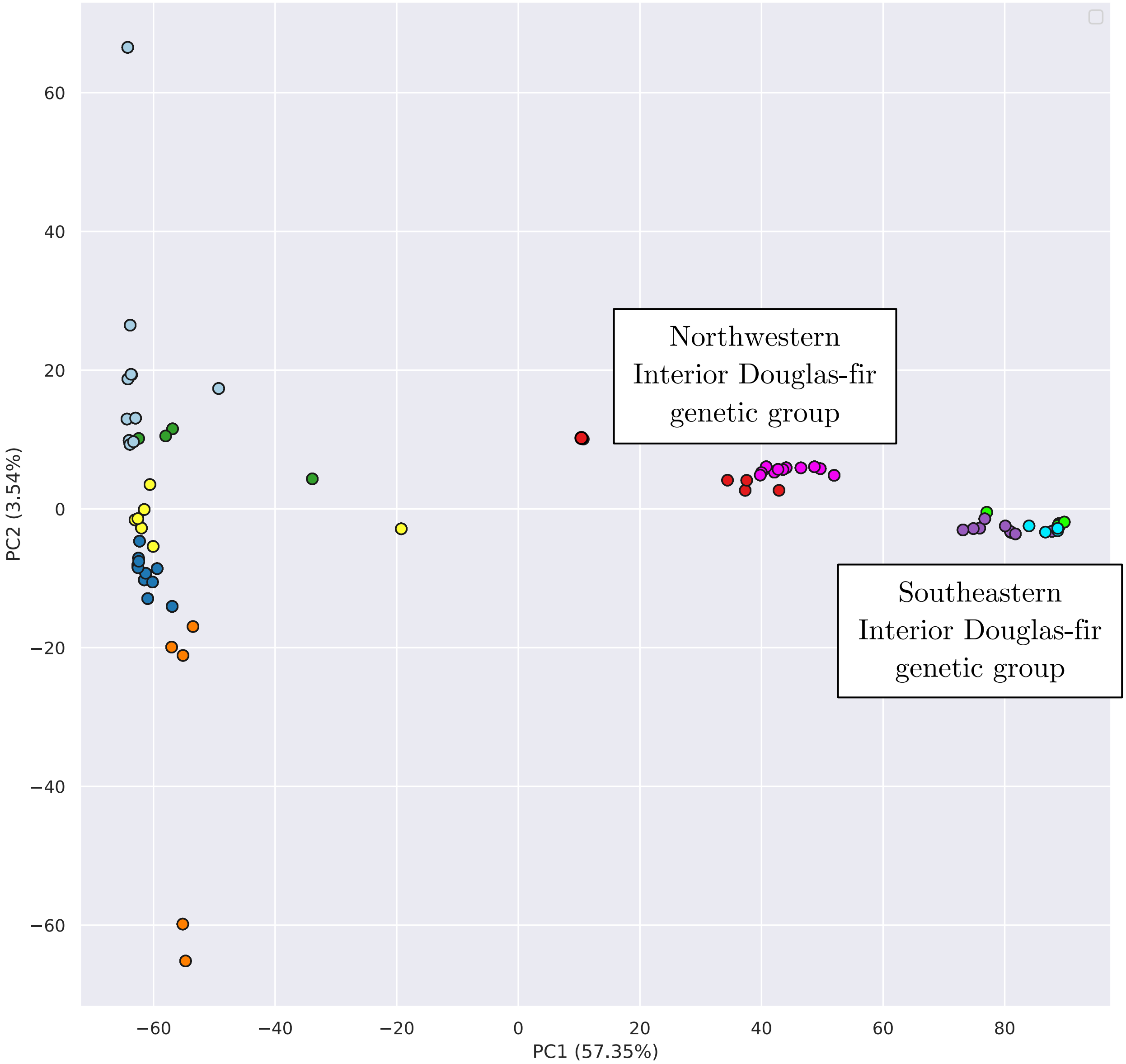


**Fig. S1** Principal components analysis (PCA) of the two varieties of Douglas-fir using SNPs called across both varieties. Populations are colored as in Fig. 1 of the main text. Code used to create this figure can be found in SN 02.12.02.

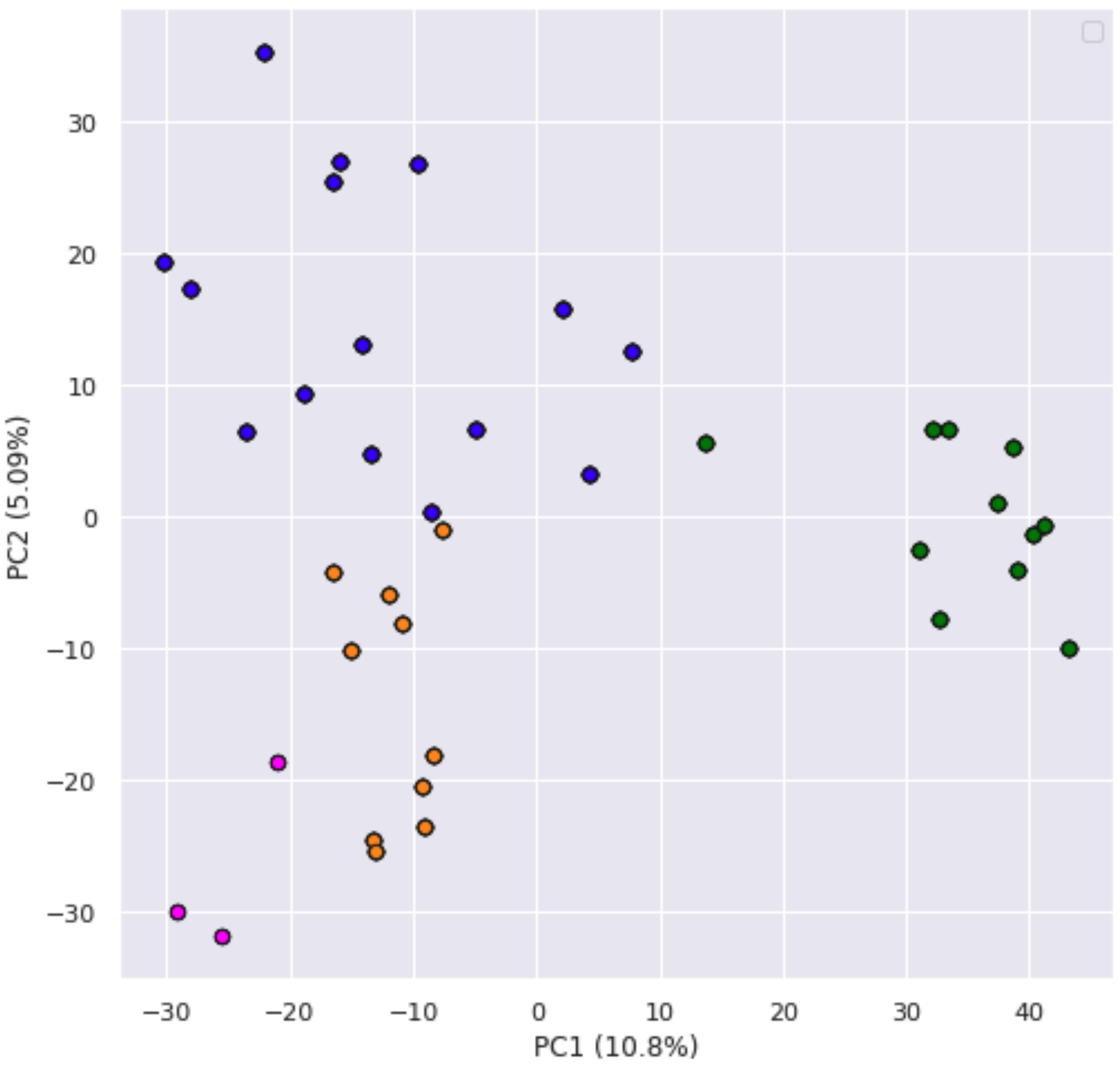


**Fig. S2** Principal components analysis (PCA) of jack pine based on SNP data. Populations are colored as in Fig. 1 of the main text. Code used to create this figure can be found in SN 02.12.02.

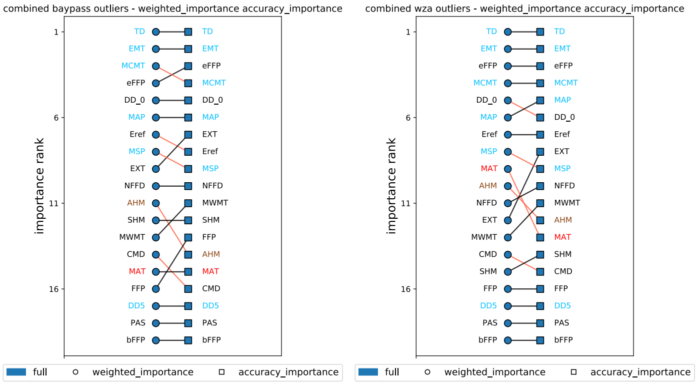
combined baypass outliers - weighted_importance accuracy_importance
importance rank
1 6 11 16
TD EMT MCMT eFFP DD_0 MAP Eref MSP EXT NFFD AHM SHM MWMT CMD MAT FFP DD5 PAS bFFP
TD EMT eFFP MCMT DD_0 MAP EXT Eref MSP NFFD MWMT SHM FFP AHM MAT CMD DD5 PAS bFFP
full weighted_importance accuracy_importance
combined wza outliers - weighted_importance accuracy_importance
importance rank
1 6 11 16
TD EMT eFFP MCMT DD_0 MAP Eref MSP MAT AHM NFFD EXT MWMT CMD SHM FFP DD5 PAS bFFP
TD EMT eFFP MCMT MAP DD_0 Eref EXT MSP NFFD MWMT AHM MAT SHM CMD FFP DD5 PAS bFFP
full weighted_importance accuracy_importance
A

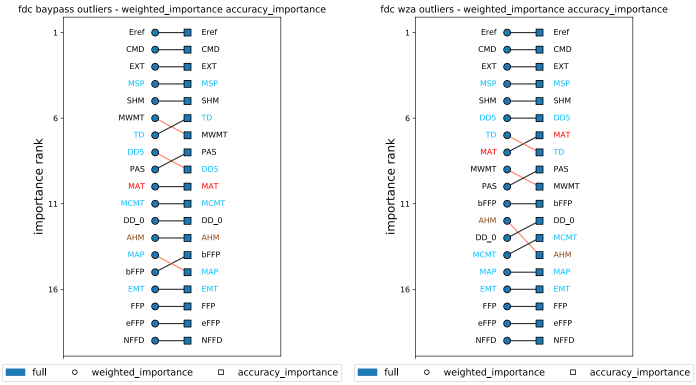
fdc baypass outliers - weighted_importance accuracy_importance
importance rank
1 6 11 16
Eref CMD EXT MSP SHM MWMT TD DD5 PAS MAT MCMT DD_0 AHM MAP bFFP EMT FFP eFFP NFFD
Eref CMD EXT MSP SHM TD MWMT PAS DD5 MAT MCMT DD_0 AHM bFFP MAP EMT FFP eFFP NFFD
full weighted_importance accuracy_importance
fdc wza outliers - weighted_importance accuracy_importance
importance rank
1 6 11 16
Eref CMD EXT MSP SHM DD5 TD MAT MWMT PAS bFFP AHM DD_0 MCMT MAP EMT FFP eFFP NFFD
Eref CMD EXT MSP SHM DD5 MAT TD PAS MWMT bFFP DD_0 MCMT AHM MAP EMT FFP eFFP NFFD
full weighted_importance accuracy_importance
B

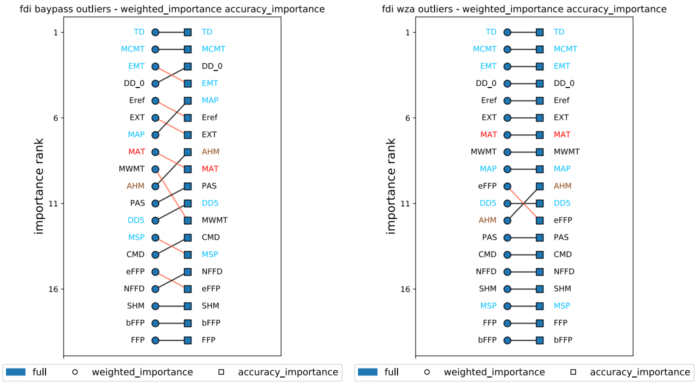
fdi baypass outliers - weighted_importance accuracy_importance
importance rank
1 6 11 16
TD MCMT EMT DD_0 Eref EXT MAP MAT MWMT AHM PAS DD5 MSP CMD eFFP NFFD SHM bFFP FFP
TD MCMT DD_0 EMT MAP Eref EXT AHM MAT PAS DD5 MWMT CMD MSP NFFD eFFP SHM bFFP FFP
full weighted_importance accuracy_importance
fdi wza outliers - weighted_importance accuracy_importance
importance rank
1 6 11 16
TD MCMT EMT DD_0 Eref EXT MAT MWMT MAP eFFP DD5 AHM PAS CMD NFFD SHM MSP FFP bFFP
TD MCMT EMT DD_0 Eref EXT MAT MWMT MAP AHM DD5 eFFP PAS CMD NFFD SHM MSP FFP bFFP
full weighted_importance accuracy_importance
C

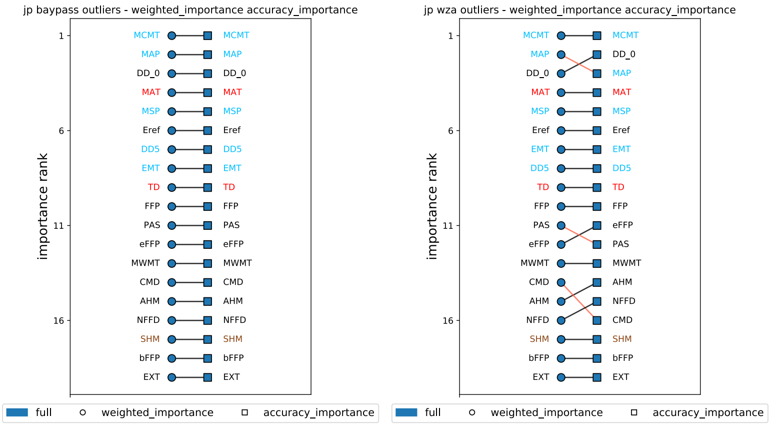
jp baypass outliers - weighted_importance accuracy_importance
importance rank
1 6 11 16
MCMT MAP DD_0 MAT MSP Eref DD5 EMT TD FFP PAS eFFP MWMT CMD AHM NFFD SHM bFFP EXT
MCMT MAP DD_0 MAT MSP Eref DD5 EMT TD FFP PAS eFFP MWMT CMD AHM NFFD SHM bFFP EXT
full weighted_importance accuracy_importance
jp wza outliers - weighted_importance accuracy_importance
importance rank
1 6 11 16
MCMT MAP DD_0 MAT MSP Eref EMT DD5 TD FFP PAS eFFP MWMT CMD AHM NFFD SHM bFFP EXT
MCMT DD_0 MAP MAT MSP Eref EMT DD5 TD FFP eFFP PAS MWMT AHM NFFD CMD SHM bFFP EXT
full weighted_importance accuracy_importance
D

**Fig. S5** Consistency of environmental accuracy and weighted importance rank output from trained models of Gradient Forests using candidate loci sets (BayPass and WZA) for A) the cross-variety model, B) FDC coastal variety, C) FDI interior variety, D) JP jack pine. Code to create these figures can be found in SN 15.13.

**Fig S6.**

**Cross-variety model – WZA**

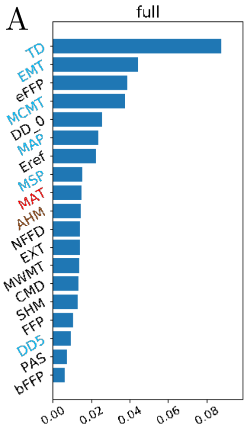


**Cross-variety model – baypass**

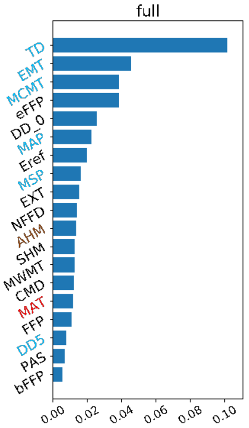

**Fig S6 (cont'd)**

**Coastal variety model – WZA**

C

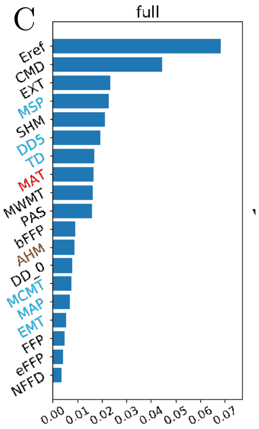


**Coastal variety model – baypass**

D

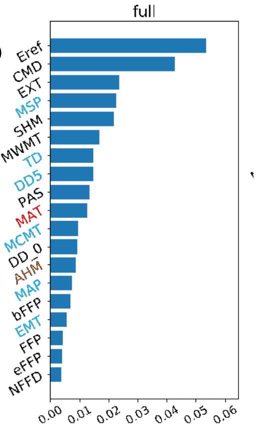

**Fig S6 (cont'd)**

**Interior variety model – WZA**

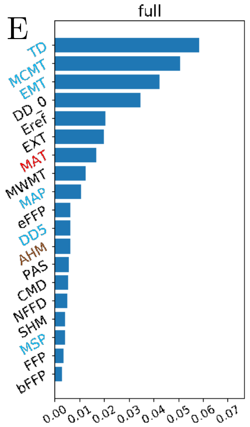


**Interior variety model – baypass**

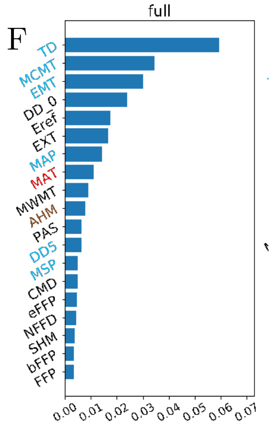

**Fig S6 (cont'd)**

**Jack pine - WZA**

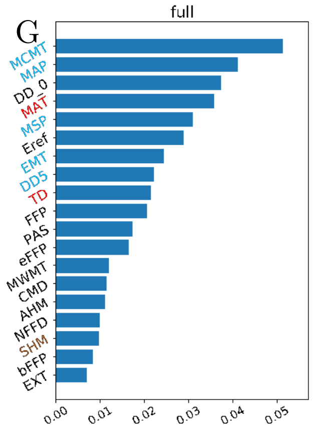


**Jack pine - baypass**

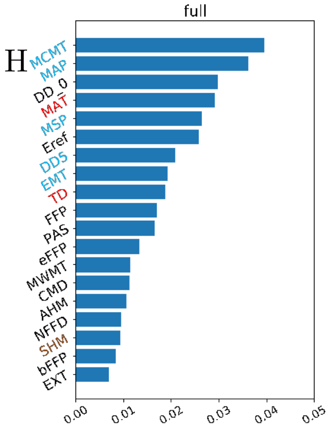


**Fig S6.** Weighted feature (climate) importance from Gradient Forest models trained using either WZA (A, C, E, G) or BayPass loci (B, D, F, H) using both varieties of Doulgas-fir (A, B), the coastal variety of Douglas-fir (C, D), the interior variety of Douglas-fir (E, F), or jack pine (G, H). Colors refer to the set of populations used: blue – all populations. Code to create this figure can be found in SN 15.13.

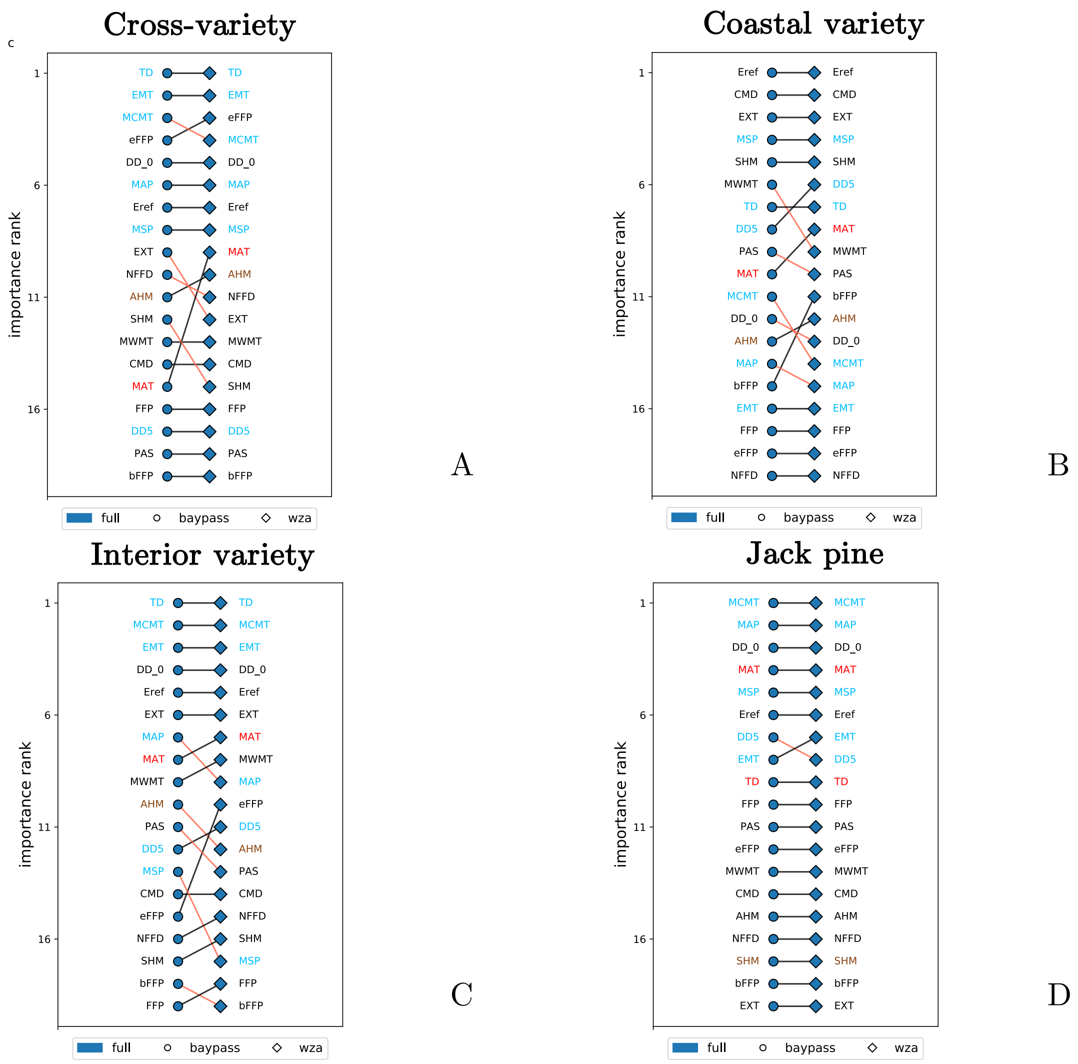


**Fig. S7** Consistency of environmental weighted importance rank across models of Gradient Forests trained using outlier marker sets from BayPass (circles) or the WZA (diamonds) and either all populations (full, blue) or *k*-fold stratified sampling (orange, green, red, and purple) for A) both Douglas-fir varieties, B) coastal Douglas-fir, C) interior Douglas-fir, and D) jack pine. Within each figure, red lines indicate negative changes in rank, while black lines indicate changes in rank ≥0. Environmental labels are colored according to membership within 1) those used in climate-based seed transfer (CBST) guidelines in British Columbia (sky blue), 2) those explaining significant variation from provenance trials (saddle brown), 3) those overlapping CBST and provenance trial environments (red), and 4) remaining environments (black). Histograms of these values can be found in Supplemental Fig. S6. Code used to create this figure can be found in SN 15.13.

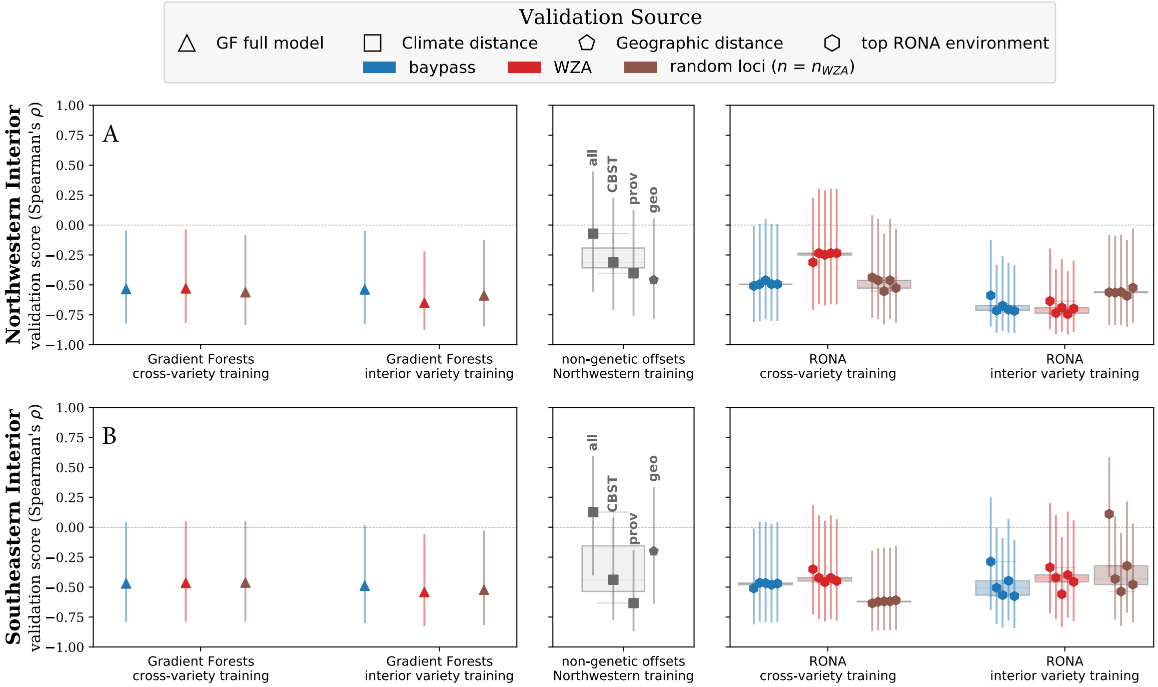


**Fig. S9** Offset validation from two-year Douglas-fir height increment phenotypes at the Vancouver common garden (see Fig. 1A) using Gradient Forests (GF), the Risk of Non-Adaptedness (RONA), and climate and geographic distances. Gradient Forests boxplots are indicative of hold-out validation from stratified sampling using out-of-bag populations (circles) while triangles indicate performance of models trained and validated using all available populations. RONA boxplots are indicative of the range of RONA validation scores given for the top five climatic variables (hexagons) that differed significantly between source (i.e., populations used in validation) and common garden variables (see Table S1). Climate distances (squares) were calculated using 1) all climate variables, or 2) those variables used for climate-based seed transfer (CBST) in British Columbia, or 3) those explaining significant variation in provenance trials. We used genetic hierarchy to assess accuracy inference using the validation statistic at the variety level for coastal and interior varieties, and at the subvariety level within the interior variety. X-axis labels indicate populations used in training sets, and the rows indicates populations used in validation sets. Vertical bars indicate standard error estimated using a Fisher transformation (see Supplemental Text S1.5). Locus counts in Extended Data Table 1. See Fig. S10 for similar validation using shoot biomass. See Supplemental Fig. S9 for all locus groups. Boxplot whiskers extend up to 1.5x the interquartile range. Code to create these figures can be found in SN 15.14.

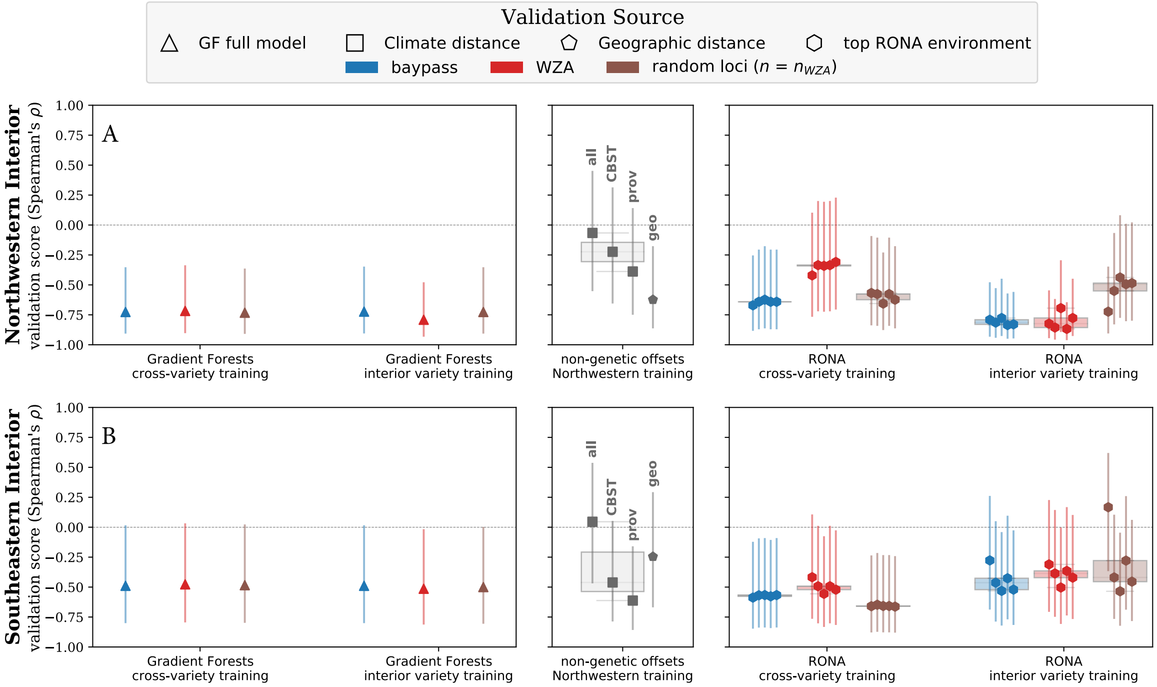


**Fig. S10** Offset validation from two-year Douglas-fir shoot biomass phenotypes at the Vancouver common garden (see Fig. 1A) using Gradient Forests (GF), the Risk of Non-Adaptedness (RONA), and climate and geographic distances. Genomic offset boxplots and shapes are shaded with respect to marker set source. Gradient Forests boxplots are indicative of hold-out validation from stratified sampling using out-of-bag populations (circles) while triangles indicate performance of models trained and validated using all available populations. RONA boxplots are indicative of the range of RONA estimates given for the top five climatic variables (hexagons) that differed significantly between source (i.e., populations used in validation) and common garden variables (see Table S1). Climate distances (squares) were calculated using 1) all climate variables, or 2) those variables used for climate-based seed transfer (CBST) in British Columbia, or 3) those explaining significant variation in provenance trials. We used genetic hierarchy to assess accuracy inference using the validation statistic at the variety level for coastal and interior varieties, and at the subvariety level within the interior variety. X-axis labels indicate populations used in training sets, and the rows indicates populations used in validation sets. Vertical bars indicate error estimated using a Fisher transformation (see Supplemental Text S1.5). Locus counts in Extended Data Table 1. See Fig. S10 for similar validation using shoot biomass. See Supplemental Fig. S9 for all locus groups. Boxplot whiskers extend up to 1.5x the interquartile range. Code to create these figures can be found in SN 15.14.

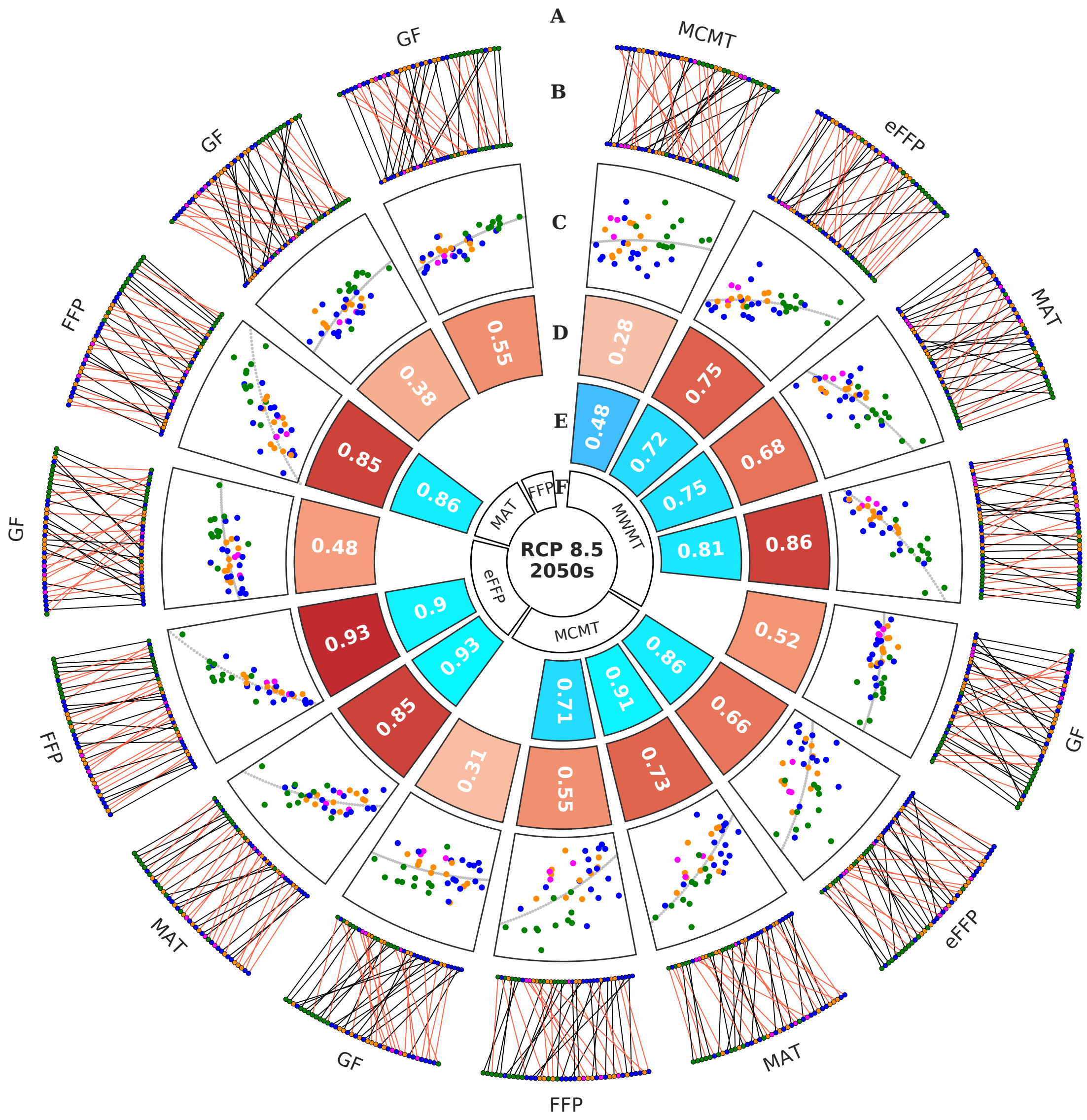


**Fig S11** Comparisons of Gradient Forest (GF) and RONA top environments for projected maladaptation to RCP 8.5 2050s for jack pine. Circos plots show pairwise comparisons (A and F) between the offset predicted from the top five environments from RONA and GF (B-D) where B* shows the changes in rank between comparisons, C** shows the linear relationship (with gray line of best fit), and D shows the Spearman's rank correlation of offset predictions. E shows the correlation between future environmental variables compared in A and F. Environments used to estimate RONA were chosen based on ranking p-values from paired *t*-tests between current and future climate across all populations that went into a given model . Populations in B, C, and G are colored as in Fig. 1. Red lines in B are indicative of negative changes in rank between F and A. Code to create this figure can be found in SN 15.17. Analogous figures created using climate models RCP4.5 2080s, RCP4.5 2050s, and RCP8.5 2080s show similar patterns and are not shown except within SN 15.17.

* outer ranks are from A, inner ranks are from F
** y-axis is from A and x-axis is from F

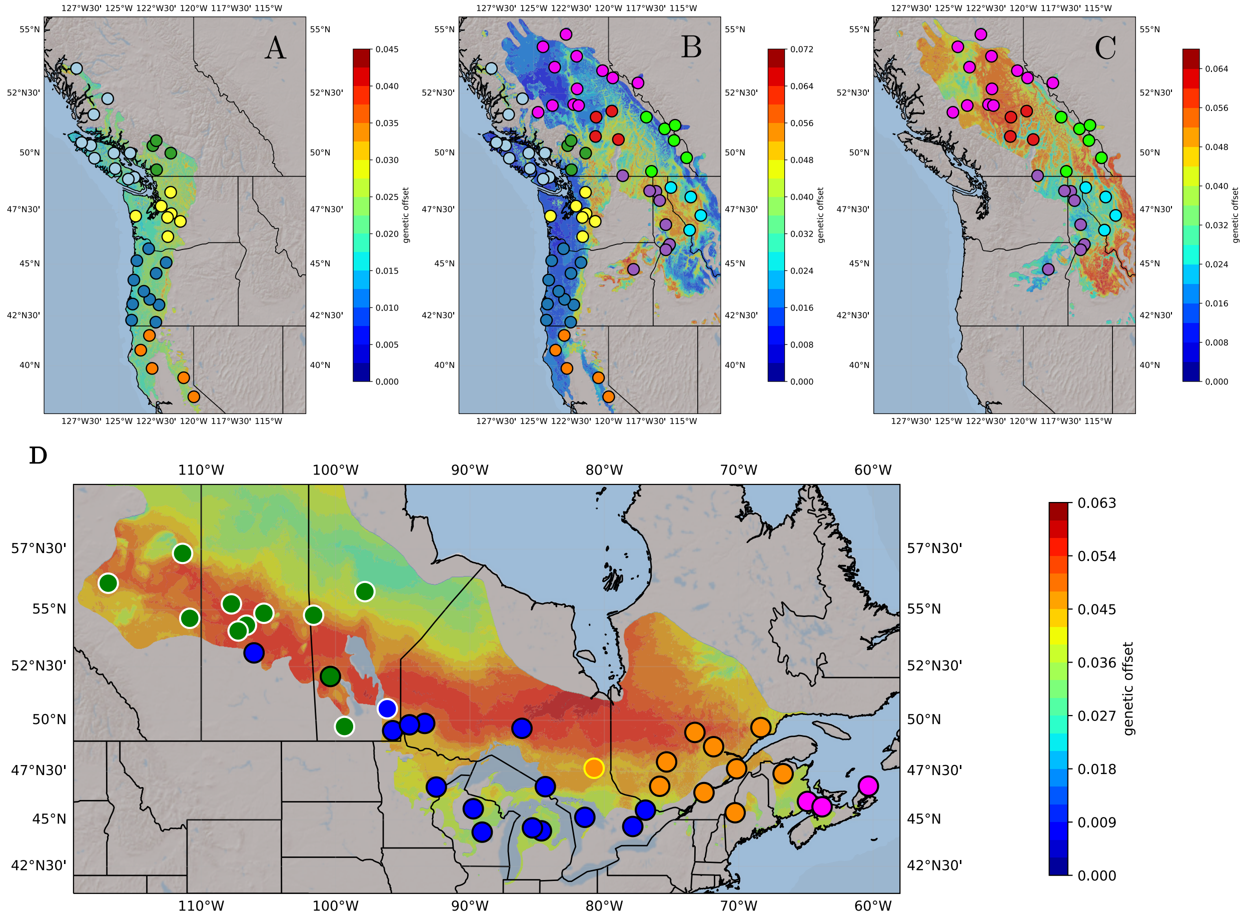


**Fig S12** Maladaptation of Douglas-fir (A-C) and jack pine (D) populations to future climate (RCP8.5 2050s) inferred from Gradient Forests. These figures are identical to maps shown in Figs. 4-5, except with the overlay of sample populations. Note color of population corresponds to Fig. 1 and does not reflect values of predicted offset. Code to create these figures can be found in SN 15.18.

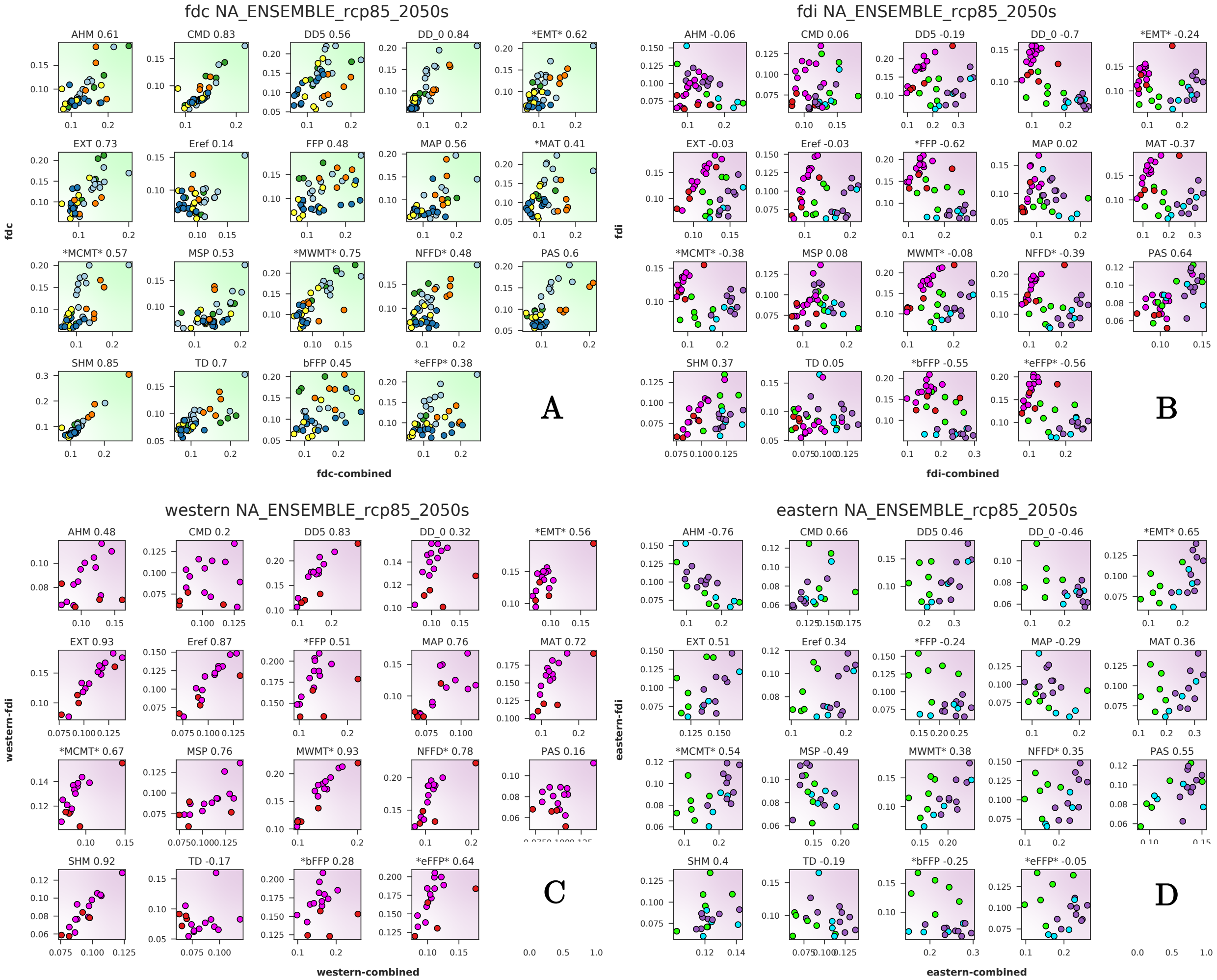


**Fig S13** Relationship between Douglas-fir RONA offset predictions to future climates (RCP 8.5 2050s) from variety-specific models and cross-variety models (A, B) or for genetic groups in the interior variety-specific models with the same genetic groups taken from the cross-variety model (C, D). Environmental names are at the top of each plot – an asterisk on the left side of an environmental name indicates that the variable was significantly different between current and future climates at the variety level; similarly an asterisk on the right side of an environmental variable indicates a significant difference in climate between current and future conditions using populations from both varieties. Background color of each figure indicates population membership to either the coastal variety (lime green) or the interior variety (purple). FDI = interior Douglas-fir; FDC = coastal Douglas-fir; eastern and western refer to the genetic groups within interior Douglas-fir; combined refers to the cross-variety model. Compare with Fig. 5 in the main text comparing similar datasets output from Gradient Forests. Code to create these figures can be found in SN 15.17.

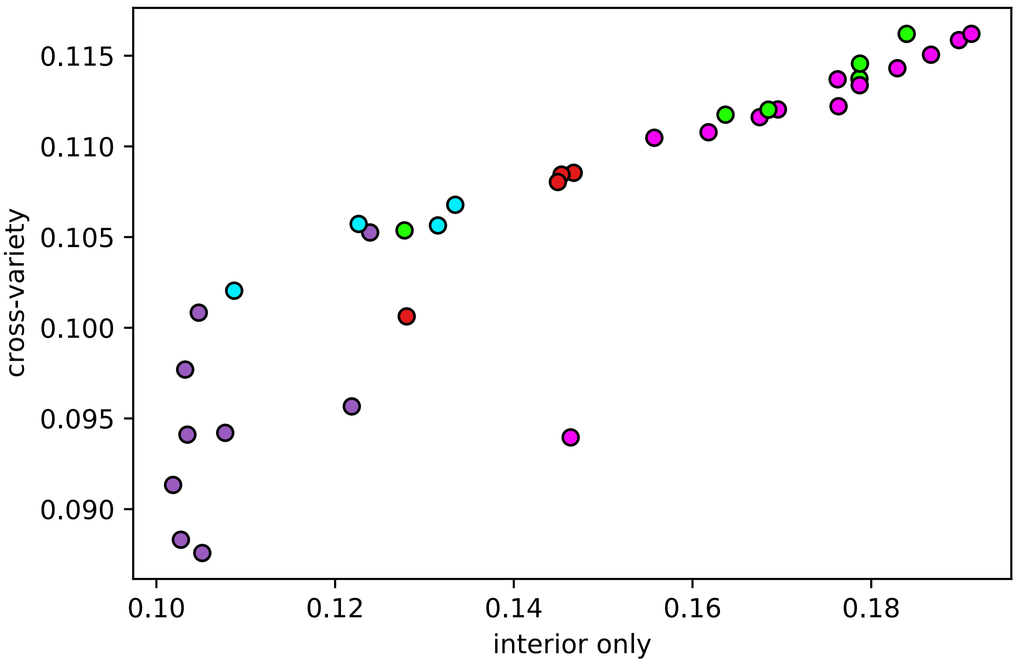


**Fig. S14** Gradient Forest (GF) model agreement as to whether the northwestern or southeastern genetic group of interior Douglas-fir is most maladapted to the climate of the Vancouver common garden used for validation. Note that the same GF models were in disagreement as to whether either genetic group was most maladapted to RCP 8.5 2050s. Models shown are the cross-variety (y-axis) and interior-only (x-axis) models of GF trained using WZA loci and all populations. Population colors are as in Fig. 1 of the main text. Code used to create this figure can be found in SN 15.20.

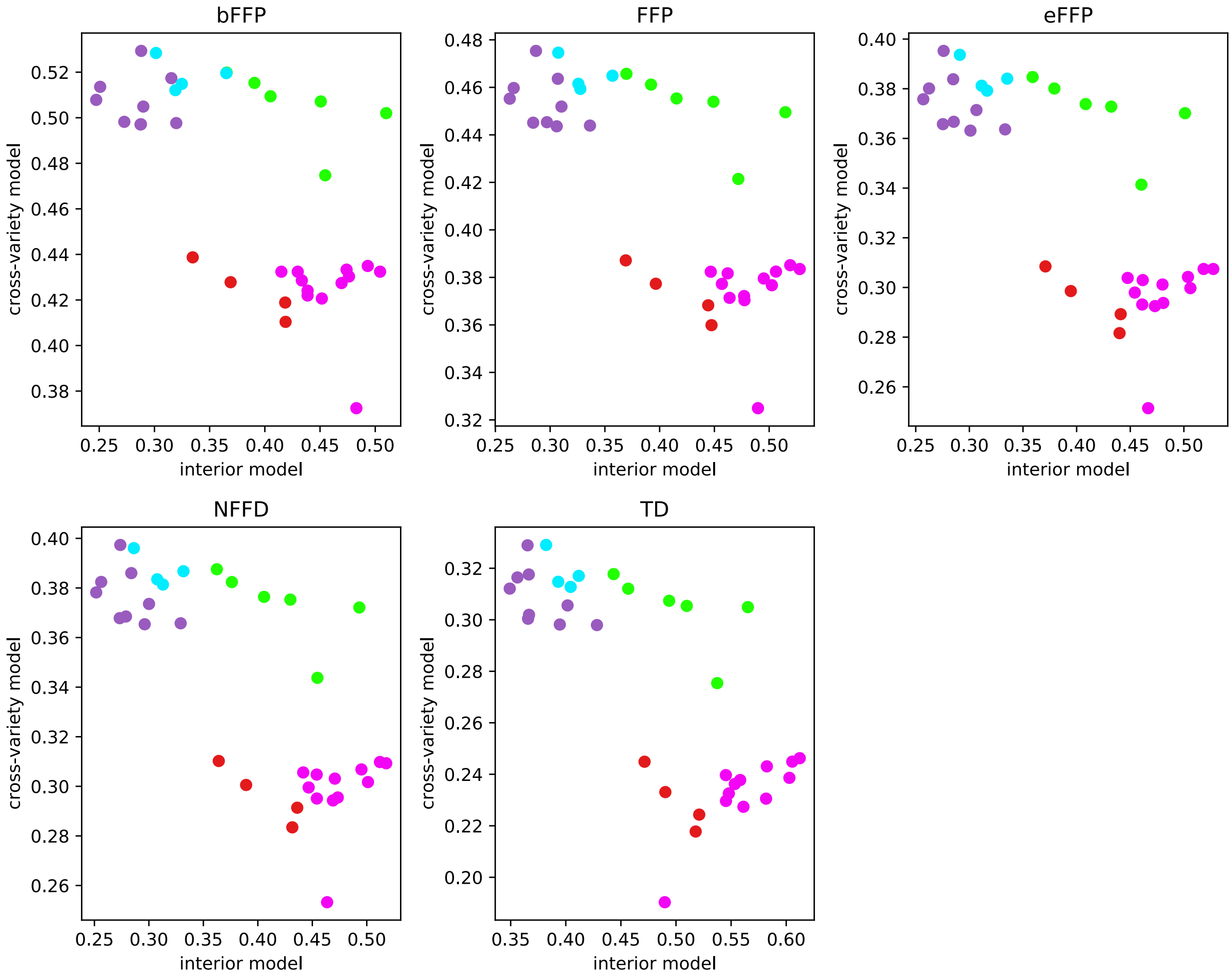


**Fig. S15** RONA model comparison of cross-variety (y-axes) and interior-only (x-axes) offset projected to the Vancouver common garden. Population colors are as in Fig. 1 of the main text. Code used to create this figure can be found in SN 15.20.

Fig S21. Comparison of Douglas-fir offset projections

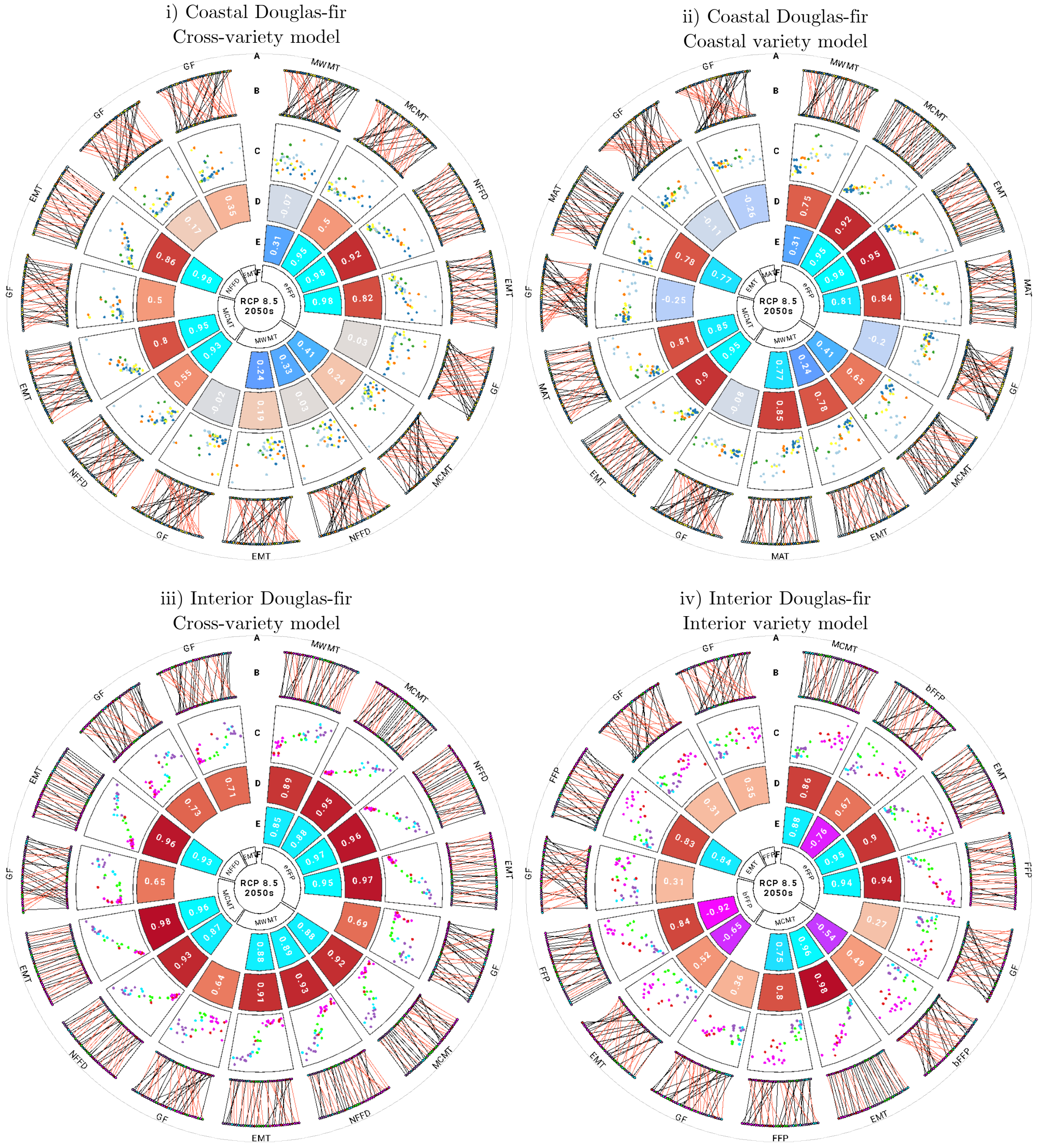

Fig S21. Comparison of Douglas-fir offset projections (cont'd)

v) Northwestern Interior Douglas-fir
Cross-variety model

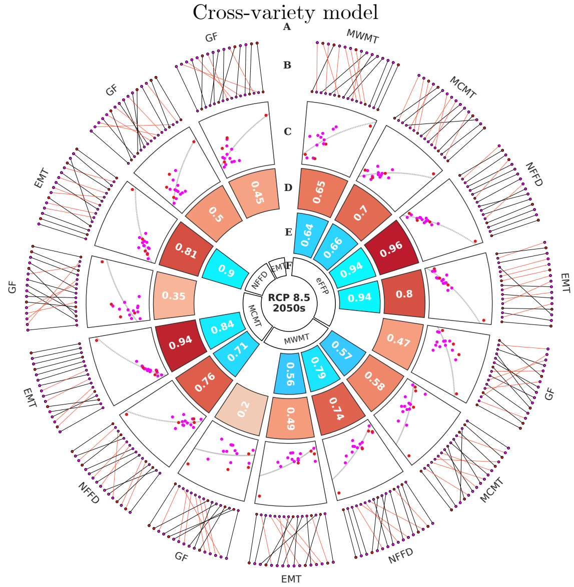


vi) Northwestern Interior Douglas-fir
Interior variety model

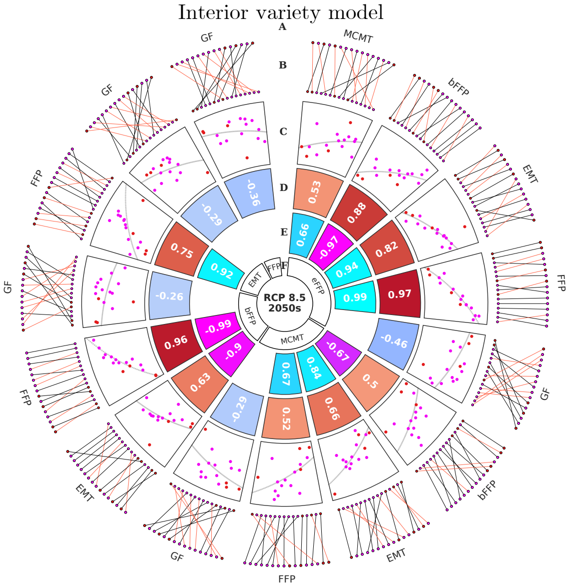


vii) Souttheastern Interior Douglas-fir
Cross-variety model

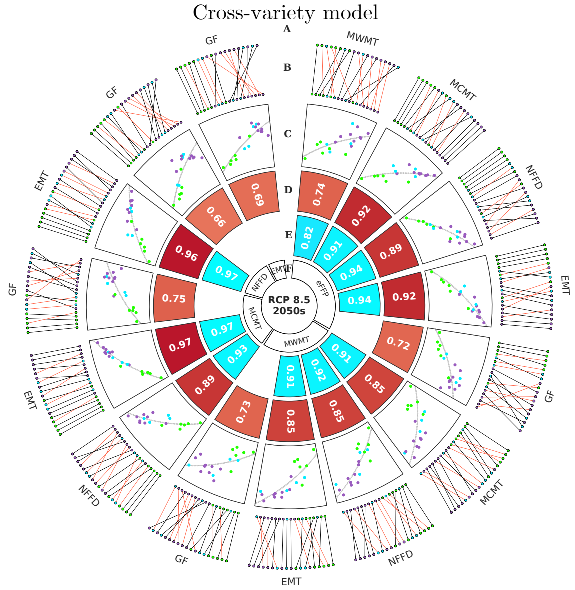


viii) Southeastern Interior Douglas-fir
Interior variety model

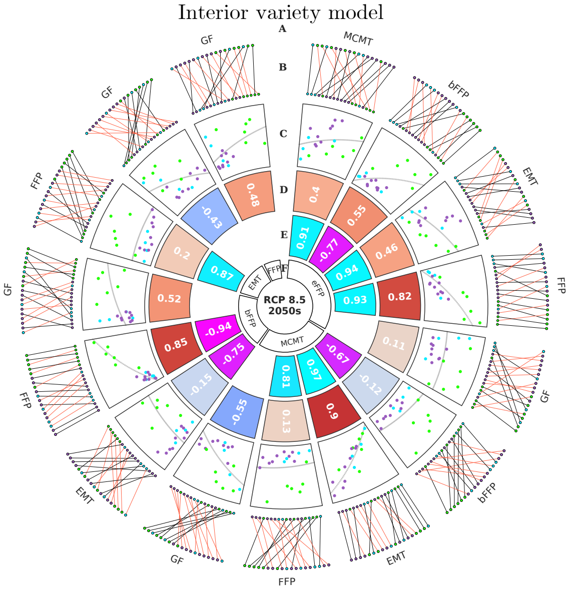

ix) Both Douglas-fir varieties
cross-variety model

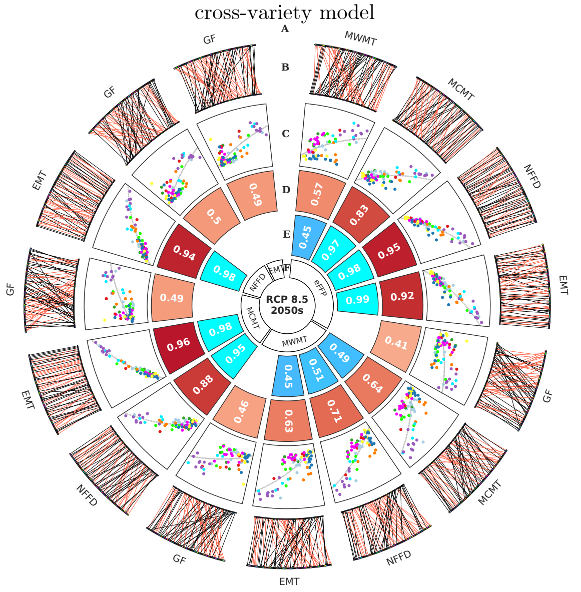


**Fig S21** Comparisons of Gradient Forest (GF) and RONA top environments for projected maladaptation to RCP 8.5 2050s (models labeled within each panel). Circos plots show pairwise comparisons (A and F) between the offset predicted from the top five environments from RONA and GF (B-D) where B* shows the changes in rank between comparisons, C** shows the linear relationship (with gray line of best fit), and D shows the Spearman's rank correlation of offset predictions. E shows the correlation between future environmental variables compared in A and F. Environments used to estimate RONA were chosen based on ranking p-values from paired *t*-tests between current and future climate across all populations that went into a given model (this is why some environments differ between cross-variety and variety-specific models). Populations in B, C, and G are colored as in Fig. 1. Red lines in B are indicative of negative changes in rank between F and A. Boxplot ranks similar to Figs. 4A, 6A-B can be found in Supplemental Fig. S22. Code to create these figures can be found in SN 15.17. Analogous figures created using climate models RCP4.5 2080s, RCP4.5 2050s, and RCP8.5 2080s show similar patterns and are not shown except within SN 15.17.

* outer ranks are from A, inner ranks are from F
** y-axis is from A and x-axis is from F

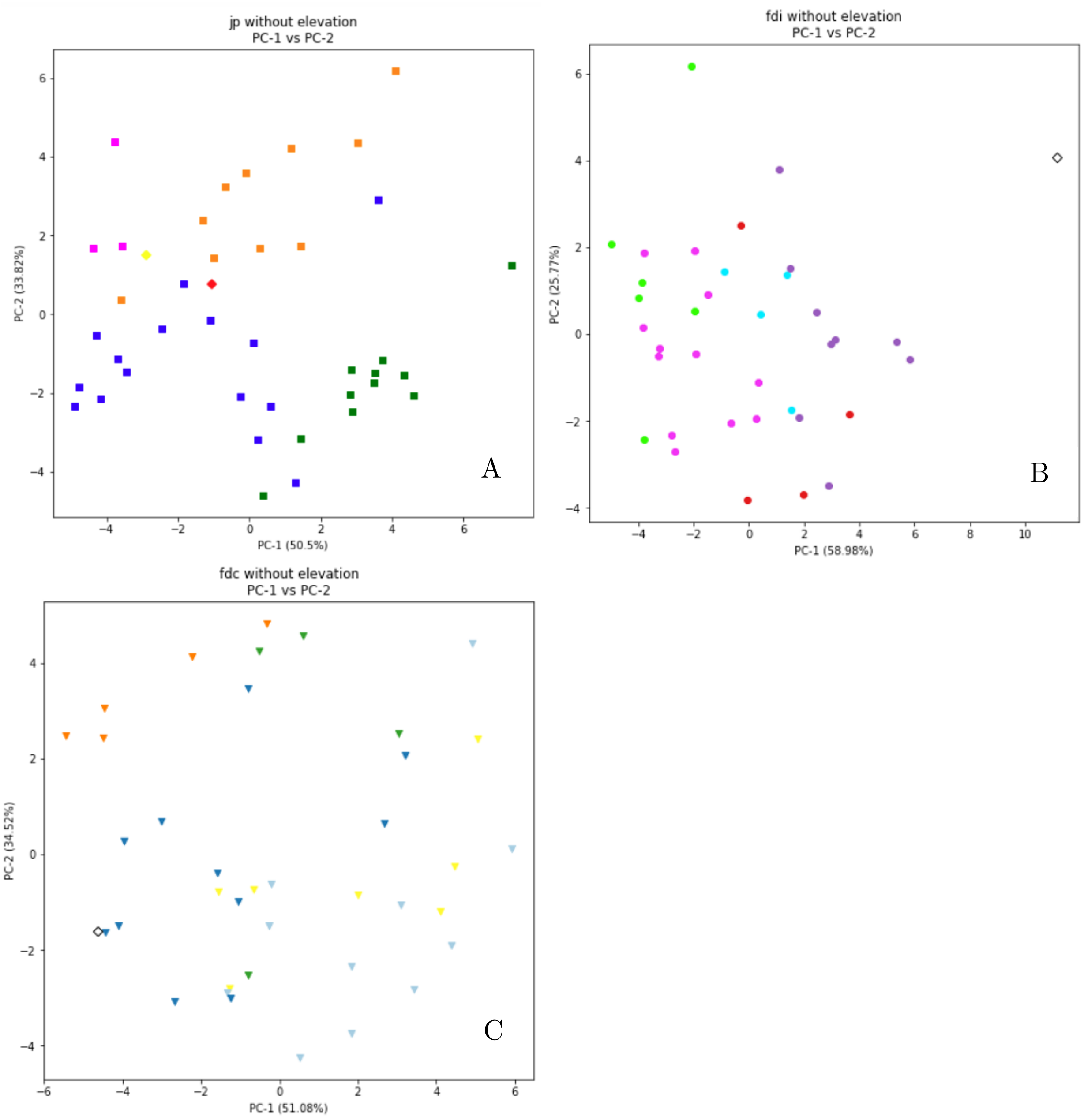


**Fig S23** Climate space as inferred from Principal Component Analysis of climate normals (1961-1990) for populations of jack pine (squares, A); coastal Douglas-fir (circles, B), and interior Douglas-fir (triangles, C) relative to common gardens used for validation (diamonds). Colors in figures are the same as the genetic groups and common gardens used in Fig. 1. Code to create these figures can be found in SN 15.19.

#### *Supplemental References*

Yeatman, C. W. 1974. The jack pine genetics program at Potawa Forest Experiment Station 1950-1970. Pp. 1–33 *in* The jack pine genetics program at Potawa Forest Experiment Station 1950-1970.

Archives of Supplemental Notebooks:

SN 15 : Lind, B.M. 2023. GitHub.com/brandonlind/offset_validation: Revision 1 release (v1.0.0). Zenodo. https://doi.org/10.5281/zenodo.7641225

SN 02 : Lind, B.M. 2023. GitHub.com/brandonlind/douglas_fir_natural_populations: Offset Revision 1 (v1.0.0). Zenodo. https://doi.org/10.5281/zenodo.8018894

SN 07 : Lind, B.M. 2023. GitHub.com/brandonlind/jack_pine_natural_populations: Offset Revision 1 (v1.0.0). Zenodo. https://doi.org/10.5281/zenodo.8018892
